# Optimal mechanical noise ensures robust vertebrate body segmentation

**DOI:** 10.1101/2021.03.27.437325

**Authors:** Arthur Boutillon, Elijah R. Shelton, Sangwoo Kim, Ben J. Gross, Ray Wu, Marie Pochitaloff, Irene Lim, Ellen M. Sletten, Otger Campàs

**Author notes:** Department of Cell and Developmental Biology, University College London, London WC1E 6BT, England, UK. Institute of Mechanical Engineering, École Polytechnique Fédérale de Lausanne (EPFL), CH-1015 Lausanne, Switzerland. These authors contributed equally (Arthur Boutillon, Elijah R. Shelton). Correspondence should be addressed to Elijah R. Shelton and/or Otger Campàs.

## Abstract

The precise and robust formation of embryonic structures is essential for the function of the organism. During somitogenesis, genetic traveling waves in the presomitic mesoderm orchestrate somite formation and the segmentation of the vertebrate body axis. While key molecular and genetic aspects of this process are known, the mechanical events required to robustly define sharp somite boundaries and physically segment the presomitic mesoderm remain unclear. Here we show that both mechanical noise in the tissue and somite boundary tension are optimized to define sharp somite boundaries and robustly segment the body axis. We find that a large, actomyosin-driven increase in boundary tension progressively straightens the nascent somite-somite boundary. While noise is typically believed to be detrimental to robustness, our results show how mechanical noise in the tissue, introduced by active tension fluctuations at cell contacts, is necessary to fully straighten somite boundaries and minimize variation across individuals. Chemical and optogenetic perturbations of both boundary tension and mechanical noise in the surrounding tissue show that zebrafish embryos optimally tune these quantities to the values necessary to ensure maximal somite boundary straightness. Altogether, these results reveal the physical mechanism of somite formation in zebrafish and uncover how optimal mechanical noise helps robustly shape embryonic structures.

## INTRODUCTION

During embryonic development, tissues and organs are formed in complex and varying environments. Ensuring that embryonic structures are robustly shaped into their functional morphologies requires molecular and/or physical mechanisms to limit variation of their morphological phenotypes^1,2^. In some cases, feedback control of signaling events helps robustly establish patterns^3,4^. In other cases, physical confinement constrains the morphogenetic outcomes to establish robust cellular packings^5,6^. However, the mechanical mechanisms that enable the precise and robust morphogenesis of embryonic structures remain elusive.

The segmentation of the vertebrate body along its anteroposterior (AP) axis into periodic, bilaterally symmetric pairs of somites, or somitogenesis (Supplementary Movie 1), is a key process in vertebrate development that lays the foundation for all skeletal muscles and vertebrae in adults^7–9^. Sharp somite boundaries enable robust embryonic development, while imprecise boundaries can lead to developmental defects, including scoliosis^10,11^. Several signaling pathways are involved in coordinating cell behaviors in space and time during the segmentation process^7,12–14^. Synchronization of genetic oscillators in neighboring cells of the presomitic mesoderm (PSM) sustain traveling waves in the tissue that emerge at the posterior end of the body^9,15–18^, travel anteriorly and eventually arrest, roughly defining the location of the new somite along the AP axis^12,16,19,20^. Eph-ephrin interactions between cells at the prospective somite boundary regulate Rho and Rac activity^21–24^, boundary cell de-adhesion and ECM deposition^25–27^. While mechanics has long been recognized as an important factor in morphogenesis and, specifically somitogenesis^28–33^, the mechanical events that help define sharp somite boundaries and ensure robust somite formation remain unclear.

## RESULTS

### Nascent somite boundaries shorten and become nearly perfectly straight during somitogenesis

Since the defining morphological event initiating the formation of a new somite in the PSM is the appearance of its posterior boundary^8^, we imaged its formation and monitored it over time by backtracking the forming somite boundary from its well-formed mature state (Fig. 1A,B; Methods; Supplementary Movie 2). The nascent posterior somite boundary appears initially contorted and sharpens over time. While the straight end-to-end distance of the nascent boundary, *L_s_*, varies only slightly during somite formation (fixed largely by the mediolateral extent of the PSM), the boundary contour length *L_c_* decreases by over 50% (Fig. 1C). This leads to the progressive straightening of the boundary (Fig. 1D and Fig. S1), characterized by the ratio *L_s_*/*L_c_*, over approximately 1 hour and ends with a nearly perfectly straight somite boundary (*L_s_*/*L_c_* ≃ 0.98). In order to temporally align (and stage) somite formation in different embryos and perform statistical analysis, we took advantage of the fact that boundary straightening during the formation of a new somite displays a sigmoid behavior (Fig. 1D), which has a well-defined timepoint (center point; t=0) characterizing the boundary straightening process (Methods). The observed nearly perfect straightening of the forming somite boundary suggests that mechanical stresses may be changing along the boundary during somite formation.

**Figure 1:**
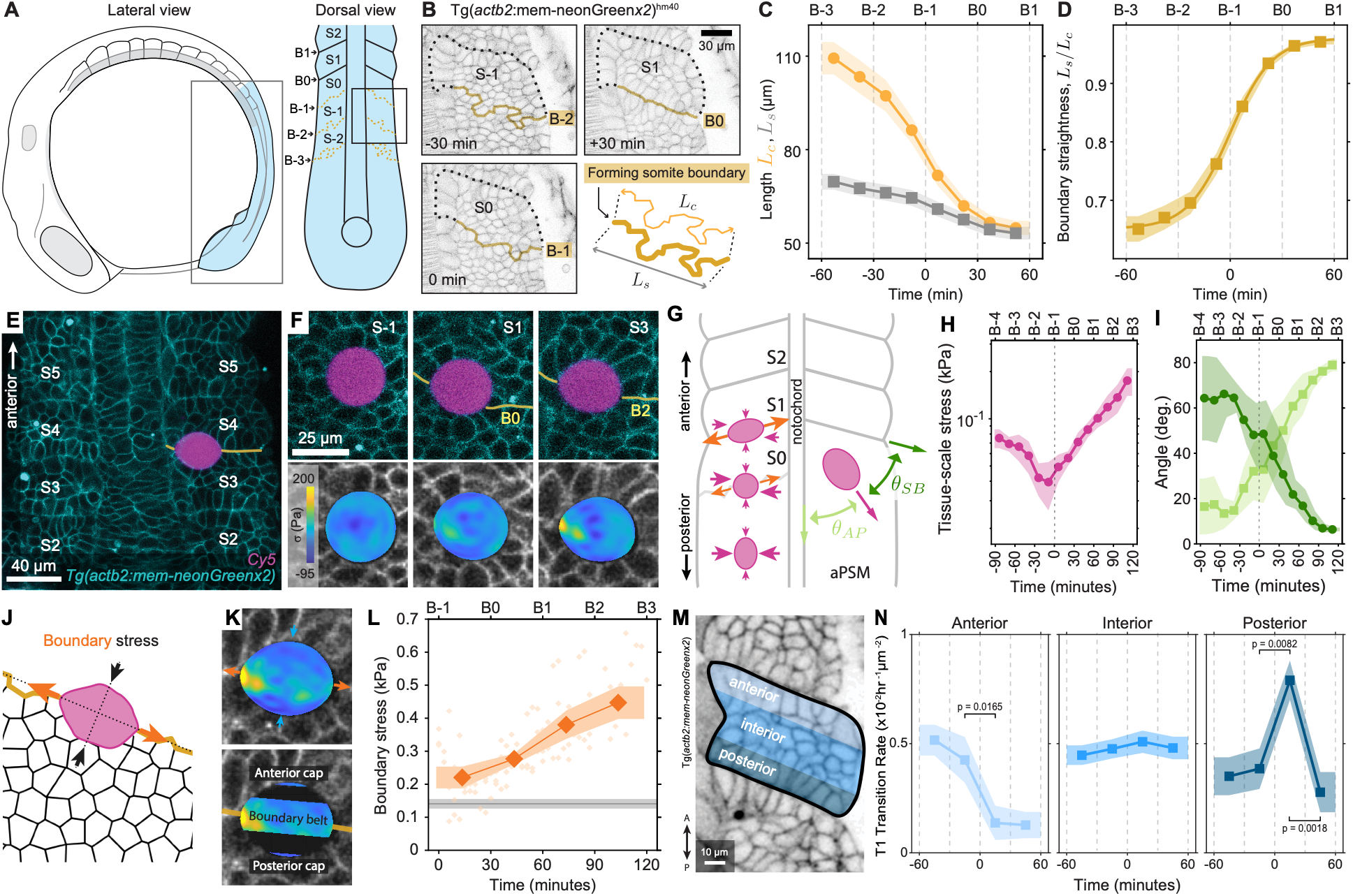
Tissue mechanical changes during somitogenesis. **A**, Sketches showing a lateral view of a 12-somite stage zebrafish embryo and a dorsal view of posterior tissues showing somites at different maturation stages (from S-2 to S2). **B**, Confocal sections of a forming somite in Tg(*actb2*:mem-neonGreen-neonGreen)^hm40^ embryos at S-1, S0 and S1, with its boundaries highlighted with dashed lines (posterior nascent boundary, yellow solid; other boundaries, black dashed). A sketch of the forming posterior boundary is shown with the contour and end-to-end lengths, *L_c_* and *L_s_*, respectively, defined. **C-D**, Temporal evolution of *L_c_* (orange; **C**), *L_s_* (gray; **C**) and the boundary straightness *L_s_/L_c_* (**D**), with sigmoid fits shown as continuous lines; *n* = 22 somites from 11 embryos. **E**, Confocal section of a droplet (magenta) at the boundary between somites S3 and S4 (membrane label, cyan; somite boundary B3 shown in yellow). **F**, Confocal sections of the temporal evolution (S-1 to S3) of a droplet at the somite boundary (top) and corresponding droplet stress maps (bottom). **G**, Sketch showing the evolution of stresses during somitogenesis (left): increasing somite boundary stresses (orange) progressively compete with ML stresses in the PSM. Definition of the angles between droplet ellipsoidal elongation and the AP (light green) and somite boundary (dark green) directions, *θ_AP_* and *θ_SB_* respectively (right). **H-I**, Time evolution of tissue-scale stresses (**H**) and of the direction of droplet elongation (**I**), quantified by *θ_AP_* (light green) and *θ_SB_* (dark green), during somitogenesis. *N* = 4 droplets from 4 embryos. **J**, Sketch of a droplet located at the forming boundary (orange line) and forces acting on it (arrows). **K**, Confocal section of a droplet (stress colormap as in **F**) at a somite boundary (membrane label) with arrows indicating anisotropic stresses, and definitions of boundary belt and anterior and posterior caps (Methods). **L**, Temporal evolution of boundary stress. Gray line indicates tissue yield stress. *n* = 3 droplet from 3 embryos. Error bands = SE. **M**, Confocal section of a somite (membrane label) highlighting the anterior boundary region, somite interior, and posterior boundary region, by a gradient in blue intensity, from light to dark, respectively. **N**, Time evolution of average T1 transition rates for anterior, interior, and posterior regions. *n* = 278 T1 transitions in 14 somites from 7 embryos.

### Somite boundaries restructure the mechanical stresses in the tissue to segment the body axis

To directly quantify the temporal evolution of mechanical stresses during somite formation, we employed magnetically-responsive oil droplets^34,35^. After injecting a fluorescently-labeled droplet in the PSM of zebrafish embryos at the 4-6 somites stage, we monitored it over time as somites formed (Fig. 1E,F; Methods; Supplementary Movies 3 and 4). We captured a 3D timelapse of each droplet, quantified its deformations using automated 3D reconstruction and analysis software^36,37^ (Fig. 1F; Methods), and obtained the mechanical stresses from these deformations after calibrating the droplet *in situ* and *in vivo* (Fig. 1G-I; Methods).

Tissue-scale stress anisotropy, obtained from the ellipsoidal mode of droplet deformation^34,37^ (Methods), changed significantly in both magnitude and orientation as somites formed (Fig. 1G-I). Before somite formation began (*t* = *−*60 min; B-3), droplets were oriented along the AP axis in the PSM due to the presence of mediolateral (ML) stress anisotropy, as previously reported^34^. As somite formation proceeded, tissue-scale anisotropic stresses substantially decreased and reached a minimum just before B-1 (*t* = 0; Fig. 1H). After B-1, the magnitude of stress anisotropy strongly increased and the droplet aligned with the direction of the future somite boundary (Fig. 1H,I), suggesting the emergence of a tension at the somite boundary that competes with the ML stresses existing in the PSM (Fig. 1G). Analysis of mechanical stress anisotropy at somite boundaries, or boundary stress anisotropy (Fig. 1J-K; Methods), showed that boundary stresses are larger than tissue-scale stresses in the PSM before somitogenesis and increase over time (Fig. 1L), indicating that maximal stresses in the tissue occur at the forming somite boundary.

The measured values of boundary stress are larger than the previously measured yield stress in the anterior PSM^34^ (Fig. 1L), suggesting that boundary stresses may drive a plastic remodeling of the tissue adjacent to the boundary. Indeed, the rate of T1 transitions in the tissue adjacent to the forming posterior somite boundary showed a transient increase (*>* 2-fold) between t=0 (B-1) and t=30 min (B0) (Fig. 1m,n and Fig. S2A-E), exactly the period when stresses along the somite boundary become dominant and boundary straightening occurs (Fig. 1D,H,L). In contrast, the T1 transition rates in the interior and anterior boundary of the forming somite remained unchanged or decreased, respectively (Fig. 1M,N). Given that plasticity is controlled by the foam-like structure of these tissues^34,38,39^, the localized increase in T1 transitions reveals a transient plastic remodeling of the tissue adjacent to the forming posterior boundary, facilitating somite shape changes.

### Actomyosin accumulation and N-cadherin reduction precede ECM assembly at the forming boundary

To understand the origin of the increasing boundary stress and of boundary straightening, we analyzed the dynamics of several molecular players known to affect tissue mechanics. Actin and myosin II enrichment at tissue boundaries has been observed in Drosophila^24,40–43^ and also in mature zebrafish somites^44,45^, but their role in the mechanics of body segmentation is unclear. Other molecular players, such as N-cadherin, integrins and ECM components (fibronectin, collagen IV), are known to change at the forming boundary and could also affect its mechanics^24^. To reveal their contributions to somite boundary straightening, we quantified their dynamics at the forming boundary (Methods). Both actin and myosin II progressively accumulate at the forming boundary as it straightens (Fig. 2A-D and Fig. S3). While actin enrichment is concomitant with boundary straightening, myosin II accumulation is delayed by a time *τ* ≃ 15 min. This actomyosin accumulation at the forming boundary correlates with the increase in boundary stress (Fig. 2E), suggesting that a progressive increase in actomyosin tension at the boundary could potentially be responsible for the reported increase in boundary stress and the straightening of the boundary. N-cadherin decreases over time as the boundary forms (Fig. 2F,G), as previously observed^27^, with the decrease closely following boundary straightening (*τ* ≃ 5 min). ECM components (Fibronectin and Collagen IV, Laminin), as well as integrin and paxillin, appear at the boundary considerably later (Fig. 2H-I; Fig. S4), with delays *τ* ranging between 35 and 55 min (Fig. 2I). To assess the role of other potential ECM components, we used Rhobo6 dye^46^ (Methods), which broadly labels glycosylated molecular components such as ECM. Rhobo6 only increased its signal at the somite boundary substantially after actomyosin accumulation and after most straightening occurred (Fig. 2H-I and Fig. S4). These results show that ECM components assemble at the somite boundary after its straightening, and therefore are not the main cause of boundary straightening.

**Figure 2:**
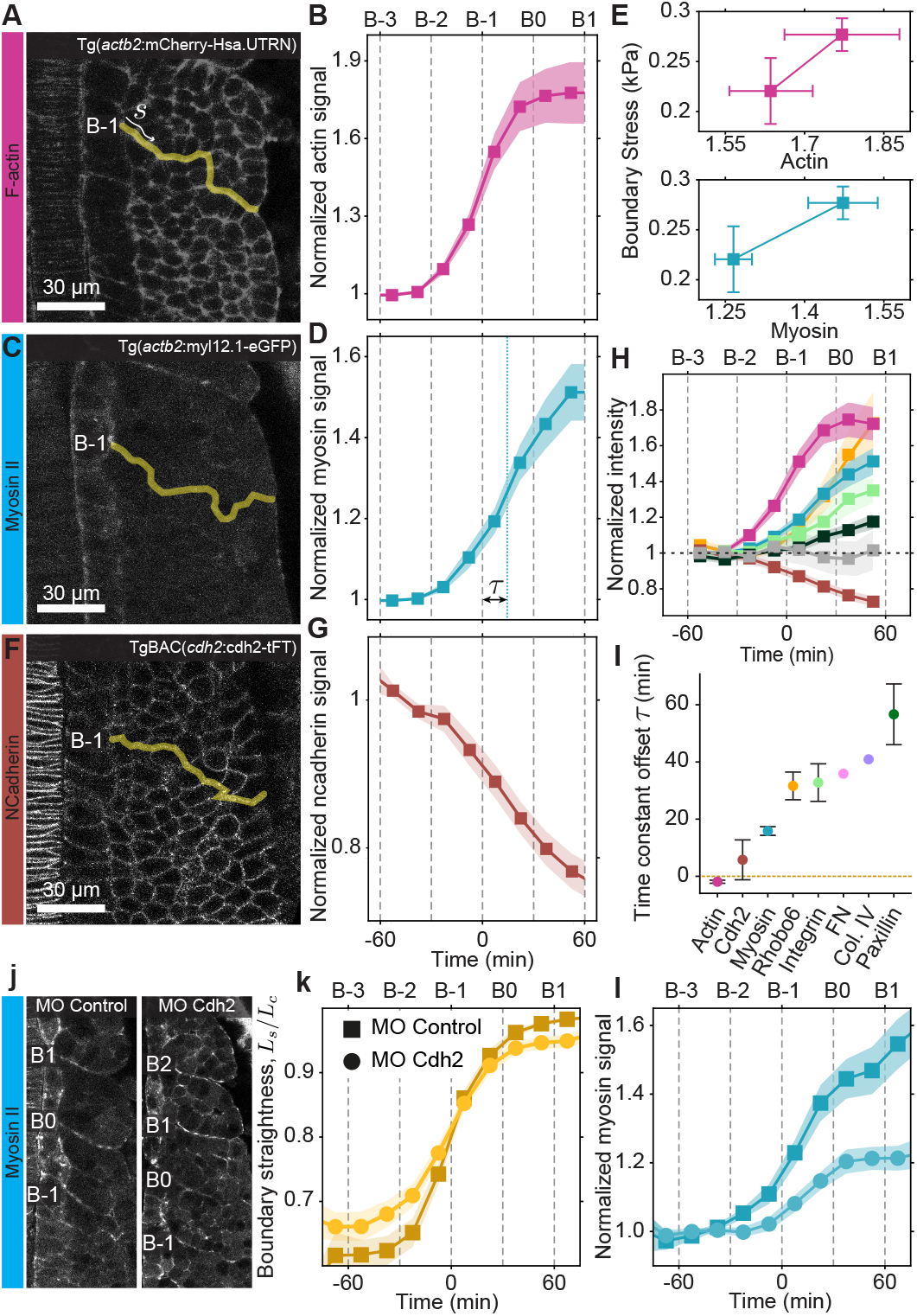
Temporal evolution of key molecular players at the forming boundary. **A-D**, **F-G**, Confocal sections of the segmenting anterior PSM showing F-actin (magenta, **A**), myosin II (cyan, **C**) and N-cadherin (brown, **F**) signals, and the respective quantifications of their temporal evolution at the forming boundary (**B**, **D**, **G**). Boundary masks are shown in yellow, and *s* is the contour distance along the boundary. n = 12, 10, 18 somites from 6, 5, 9 embryos, respectively. **E**, Relations between boundary stress (Fig. 1L) and average F-actin and myosin II densities at the somite boundary (**B**, **D**). **H**, Temporal evolution of average boundary densities for F-actin (magenta), N-cadherin (brown), myosin (blue), Rhobo6 (orange), integrin α5 (light green), paxillin (dark green) and laminin α1 (grey). n=12, 18, 10, 20, 14, 15, 7, 10 somites from 6, 9, 5, 5, 4, 6, 2, 5 embryos, respectively. **I**, Temporal shift *τ* of the sigmoidal fit of the average protein density at the forming boundary. For Fibronectin (FN, pink) and Collagen IV (Col. IV, lilac), n=209, 199 somites in 31, 27 embryos, respectively (Methods). **J**, Confocal sections of the segmenting anterior PSM showing myosin II in Tg(*actb2*:myl12.1-eGFP) embryos injected with control morpholino (MO Control) or morpholino against N-cadherin (MO Cdh2). **K-L**, Temporal evolution of boundary straightness *L_s_*/*L_c_* (**K**) and myosin II average boundary density (**L**) in embryos injected with MO Control (square dots) or MO Cdh2 (round dots). n=21, 19 from 8, 8 embryos, respectively. Error bands = SE.

The observed decrease in N-cadherin (Cdh2) adhesion at the boundary could also contribute to its straightening. However, somite formation occurs in the absence of N-cadherin adhesion in (parachute) mutants^47^ and in embryos with reduced N-cadherin expression^27^ (Methods; Fig. S5), indicating that N-cadherin is not essential for segmentation. Indeed, both somite formation (Fig. 2J) and boundary straightening (Fig. 2K) proceed normally when N-cadherin or paraxial protocadherin (Papc; Fig. S5) levels are decreased using morpholinos that recapitulate the mutant phenotype^47^ (Methods), with myosin II still accumulating at the forming boundary (Fig. 2L and Fig. S5). These results show that neither N-cadherin nor Papc drive boundary straightening and segmentation, and highlight actomyosin tension as the most likely force generating mechanism.

### Increasing actomyosin-generated tension at the forming boundary drives its straightening

Our observations of increasing myosin II and actin levels at the nascent somite boundary, concomitant with boundary straightening and increasing boundary stresses, strongly suggest that an increase in actomyosin-generated tension at the somite boundary mechanically drives segmentation events. To determine if this is indeed the case, we performed dose-dependent perturbations of myosin II activity using blebbistatin, and monitored changes in boundary straightness *L_s_*/*L_c_*, lateral boundary angle *θ* (Fig. 3A), as well as tensions at the somite boundary and cell junctions by laser ablation (Fig. 3B; Methods). While junction recoil after ablation was insensitive to changes in myosin II activity for cells in the PSM, cell junctions along the somite boundary displayed a strong suppression in their recoil velocity that depended on blebbistatin concentration (Fig. 3B), indicating the existence of actomyosin-dependent tension at the forming somite boundary. Moreover, both the boundary straightness and the lateral boundary angle between somites were affected as expected for a decrease in boundary tension in the presence of blebbistatin (Fig. 3C,D), with the maximal boundary straightness and the minimal contact angle being significantly reduced and increased, respectively, by increasing blebbistatin concentrations (Fig. 3E,F). Tissue-scale 3D ablation through the forming somite boundary showed a transient arrest in boundary formation, as measured in the dynamics of the lateral boundary angle, until the tissue healed (Fig. S6). Finally, high blebbistatin concentrations arrested boundary formation (Fig. 3G). Altogether, these results show that actomyosin contractility at the forming somite boundary generates the mechanical stresses (tension) driving boundary straightening and segmentation.

**Figure 3:**
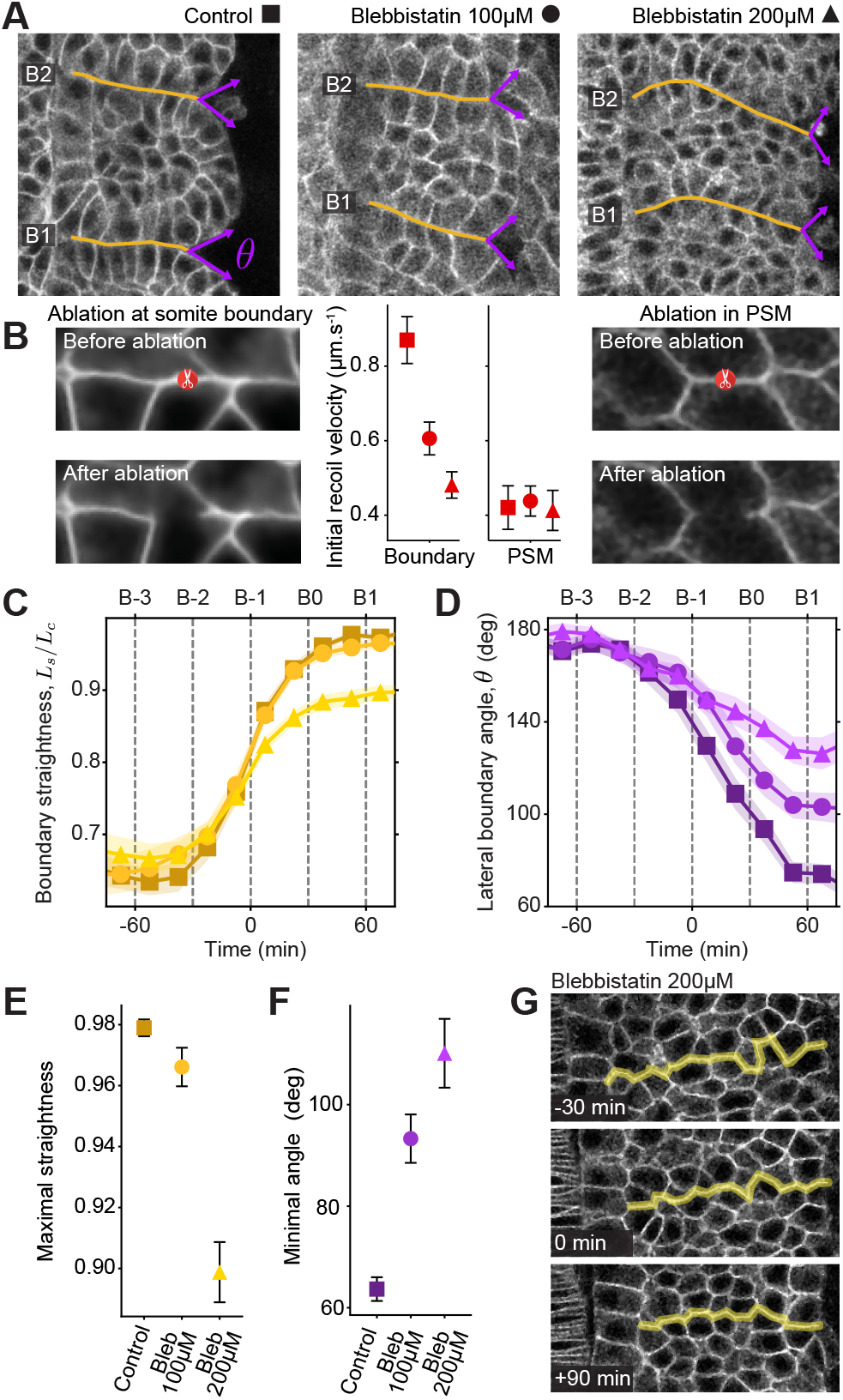
Role of myosin II activity in segmentation. **A**, Confocal sections of B1 and B2 somite boundary in Tg(actb:hRasCT-GFP) embryos with wild type phenotype (control) and incubated in 100µM or 200µM of blebbistatin. Highlighted boundaries (yellow lines) and lateral boundary angle (purple arrows) are shown. **B**, Representative confocal sections of cell junctions at the somite boundary and away from it in the PSM, before and after laser ablation (red scissors). Initial recoil velocity after laser ablation of individual cell junctions in the PSM and at the forming boundary for each condition. n=135 & 98, 152 & 103, 184 & 126 ablated junctions from 37 & 34, 35 & 30, 42 & 42 embryos for boundary and PSM in control (square), blebbistatin 100µM (circle) and blebbistatin 200µM (triangle) conditions respectively. **C-F**, Temporal evolution of boundary straightness *L_s_*/*L_c_* (**C**) and lateral boundary angle *θ* (**D**) for each condition. Maximal straightness (**E**) and minimal lateral angle (**F**) reached after straightening at B1. n=20, 22, 26 somites from 10, 9, 9 embryos for control (square), blebbistatin 100µM (circle) and blebbistatin 200µM (triangle) conditions, respectively conditions respectively. **G**, Confocal sections of a forming somite boundary (yellow) in a Tg(actb:hRasCT-GFP) embryo incubated in 200µM blebbistatin at B-2, B-1 and B2. Boundary straightening is arrested and no more somites form afterwards. Point shapes in (**B-E**) correspond to conditions as indicated in (**A**). Error bands = SE.

### Increase in boundary tension is not sufficient to robustly define sharp somite boundaries

To understand if an actomyosin-driven tension increase at the forming somite boundary was sufficient to explain segmentation, we performed simulations of the mechanics of somitogenesis using an active foam description^39^ (Methods). Starting from a population of cells in a rectangular geometry (Fig. 4A), we defined different cell-cell contacts depending on whether they are located at the lateral boundary, at a somite boundary (heterotypic contacts) or in neither of those (homotypic contacts). To computationally account for our observations that average cell-scale stresses do not change in time^34^ (Fig. S7A-B; Methods) and that boundary stresses increase, we set the average value of homotypic tensions to a constant *T*_0_, and made the average tensions at heterotypic cell-cell contacts, *T_H_*, undergo sequential ramp ups (from *T*_0_ at B-3 to a maximal value *T_M_* at B1) that follow the observed increase in actomyosin at the somite boundary (Fig. 4B,C; Methods), including the observed delay in myosin II increase (Fig. 2D) as it can affect the dynamics of straightening (Fig. S7D). Beyond their average value, it is known that cell-cell contact tensions fluctuate and drive junctional length fluctuations^34^, which introduce active mechanical noise in the tissue (Fig. 4D). Consequently, we simulated tension fluctuations of constant amplitude Δ*T* and characteristic persistence time of 90s, as observed experimentally^34,38^. Finally, we set the value of the tension at the lateral boundary to a constant *T_L_* because measurements of F-actin and myosin II at the lateral boundary show no changes over time (Fig. S7E-F).

**Figure 4:**
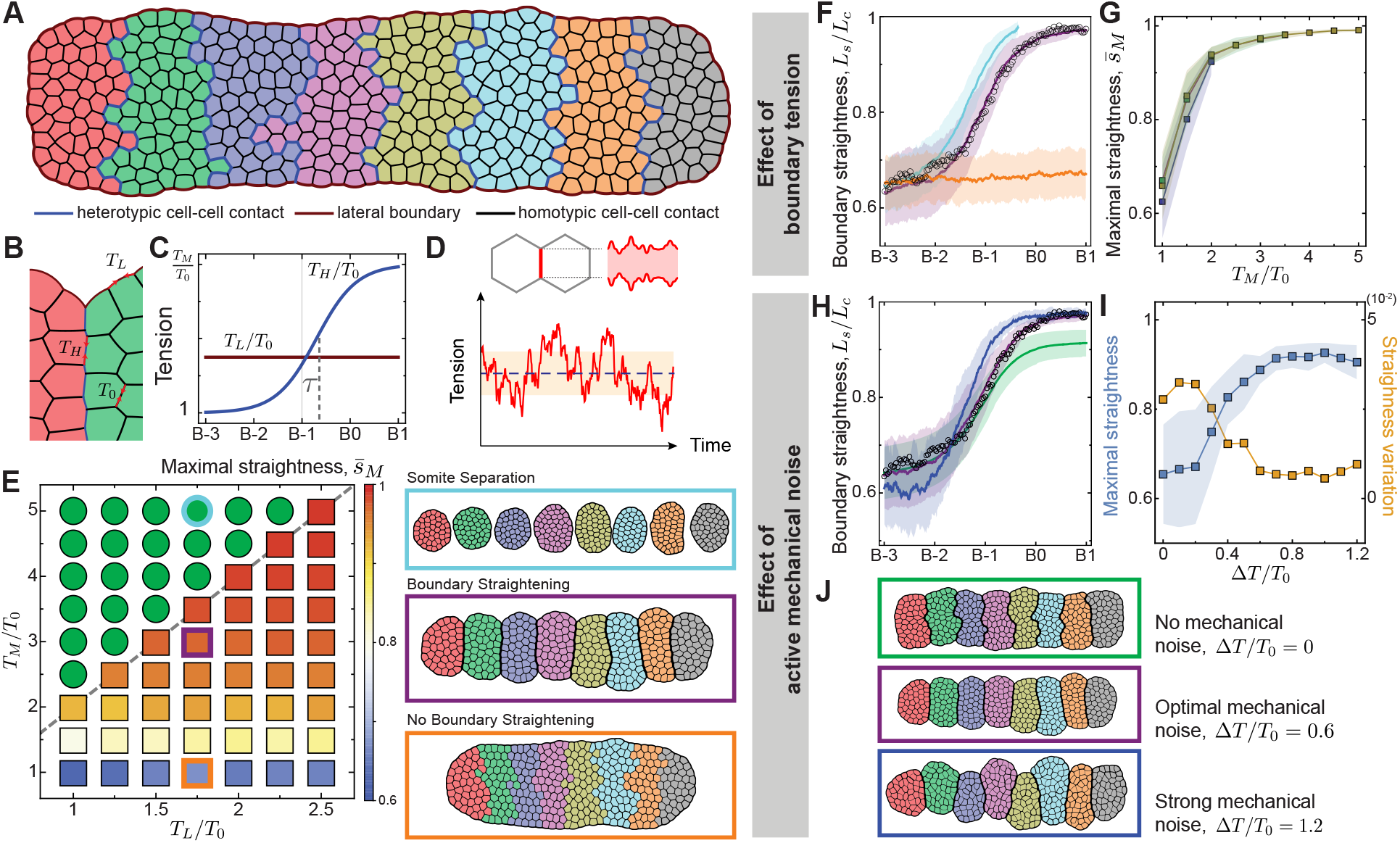
Simulations of somite formation. **A-B**, Initial state of somite formation simulation with prospective somites uniquely colored. Heterotypic (blue) and homotypic (black) cell-cell contacts, as well as the lateral boundary (maroon), are indicated. **B**, Schematic diagram showing tensions at lateral boundaries (*T_L_*), heterotypic contacts (*T_H_*), and homotypic contacts (*T*_0_). **C**, Imposed time evolution of the normalized heterotypic and lateral tensions, *T_H_*/*T*_0_ and *T_L_*/*T*_0_, respectively, with *T_H_* saturating to a maximal value *T_M_*. **D**, Example of fluctuating tensions at cell junctions in the simulations. **E**, Somitogenesis phase diagram, showing how somite formation depends on *T_L_*/*T*_0_ and *T_M_*/*T*_0_ (*ΔT*/*T*_0_ = 0.6) and snapshots of system configurations for different regimes (right). **F**, Simulated time evolution of boundary straightness *L_s_*/*L_c_* for different values of *T_M_*/*T*_0_ (same color code as in **E**; fixed *ΔT*/*T*_0_= 0.6). Measured evolution of boundary straightness is also shown (circles). **G**, Maximal boundary straightness (at B1, *L_s_*/*L_c_* (B1)) as a function of *T_M_*/*T*_0_ for different *T_L_*/*T*_0_ = 1,1.75, 2.5 (blue, green, and dark yellow, respectively; fixed *ΔT*/*T*_0_= 0.6). **H-J**, Simulated time evolution of boundary straightness *L_s_*/*L_c_* (**H**; same color code as in **J**: *ΔT*/*T*_0_ = 0 (green), 0.6 (purple), 1.2 (blue); *T_M_*/*T*_0_ = 3 and *T_L_*/*T*_0_ = 1.75; measured evolution of boundary straightness is shown (circles)), maximal boundary straightness and standard deviation of maximal boundary straightness (**i**, blue and orange, respectively; *T_M_*/*T*_0_ = 3 and *T_L_*/*T*_0_ = 1.75) for varying tension fluctuations *ΔT*/*T*_0_. Error band = SD in **F-I**.**J**, Final configurations of simulated somite formation for vanishing, optimal and strong fluctuations.

Values of the maximal heterotypic tension, *T_M_*, larger than twice the value of the lateral boundary tension, *T_L_*, namely *T_M_* > 2*T_L_*, led to complete somite separation (Fig. 4E). Below this threshold (*T_M_* < 2*T_L_*), adjacent somites shared a boundary of finite length (Fig. 4E; Supplementary Movie 5), with average values of its maximal straightness, *s̅_M_*, increasing for larger heterotypic tensions at the somite boundary (*T_M_*/*T*_0_), irrespective of the value of the tension *T_L_* at the lateral boundary (Fig. 4F,G). Increasing heterotypic tension up to 3 times homotypic tensions (*T_M_*/*T*_0_ = 3) efficiently straightened the somite boundary, but not much more above this value (Fig. 4G).

Beyond the heterotypic (boundary) tension *T_M_*, our simulations predicted that boundary straightness also depended on the magnitude of tension fluctuations Δ*T*/*T*_0_. The somite boundary could not straighten to experimentally observed values in the absence of active mechanical noise (Δ*T*/*T*_0_ = 0; Fig. 4H-J), as tension fluctuations facilitate the cell rearrangements necessary for boundary straightening^39^. Increasing the magnitude of tension fluctuations up to approximately Δ*T*/*T*_0_ ≃ 0.6 led to straighter somite boundaries (Fig. 4H-I), but not beyond this point. Moreover, the statistical variability in boundary straightness was maximal for vanishing tension fluctuations (Fig. 4I), indicating large deviations from the straight boundary, and decreased with increasing fluctuations up to about Δ*T*/*T*_0_ ≃ 0.6, defining the sharpest possible somite boundary with minimal variation at this point (Fig. 4J). In contrast, deformities in somite shape (somite angle) and deviations from the midline (positional deviations) showed a minimum at Δ*T*/*T*_0_ ≃ 0.7 (Fig. S8A). These predictions suggest the existence of an optimal value of tension fluctuations (Δ*T*/*T*_0_ ≃ 0.6) leading to a maximally sharp boundary with minimal variation (Fig. 4J).

### Optimal values of both boundary tension and active mechanical noise are necessary to define sharp somite boundaries

To test these predictions, it is necessary to change both the magnitude of maximal tension at the somite boundary (*T_M_*) and the magnitude of tension fluctuations (Δ*T*/*T*_0_). Decreasing myosin II activity with blebbistatin lowered the somite boundary (heterotypic) tension *T_M_* without affecting the average homotypic tensions *T*_0_ (Fig. 3B), and led to a dose-dependent decrease in boundary straightness and increase in the lateral boundary angle (Fig. 3C,D), in agreement with our predictions (Fig. 4F,G). Measurements of junctional length fluctuations showed that blebbistatin also affected the magnitude of active tension fluctuations, causing a dose-dependent reduction in cell-cell contact length fluctuations 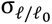 for increasing blebbistatin concentrations (Fig. 5A,B; Fig. S9). Using the relation between Δ*T*/*T*_0_ and 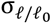 from the simulations (Fig. 5C), we mapped the experimentally measured values of 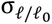 to Δ*T*/*T*_0_, providing a quantitative assessment of the dependence of Δ*T*/*T*_0_ on blebbistatin concentration.

**Figure 5:**
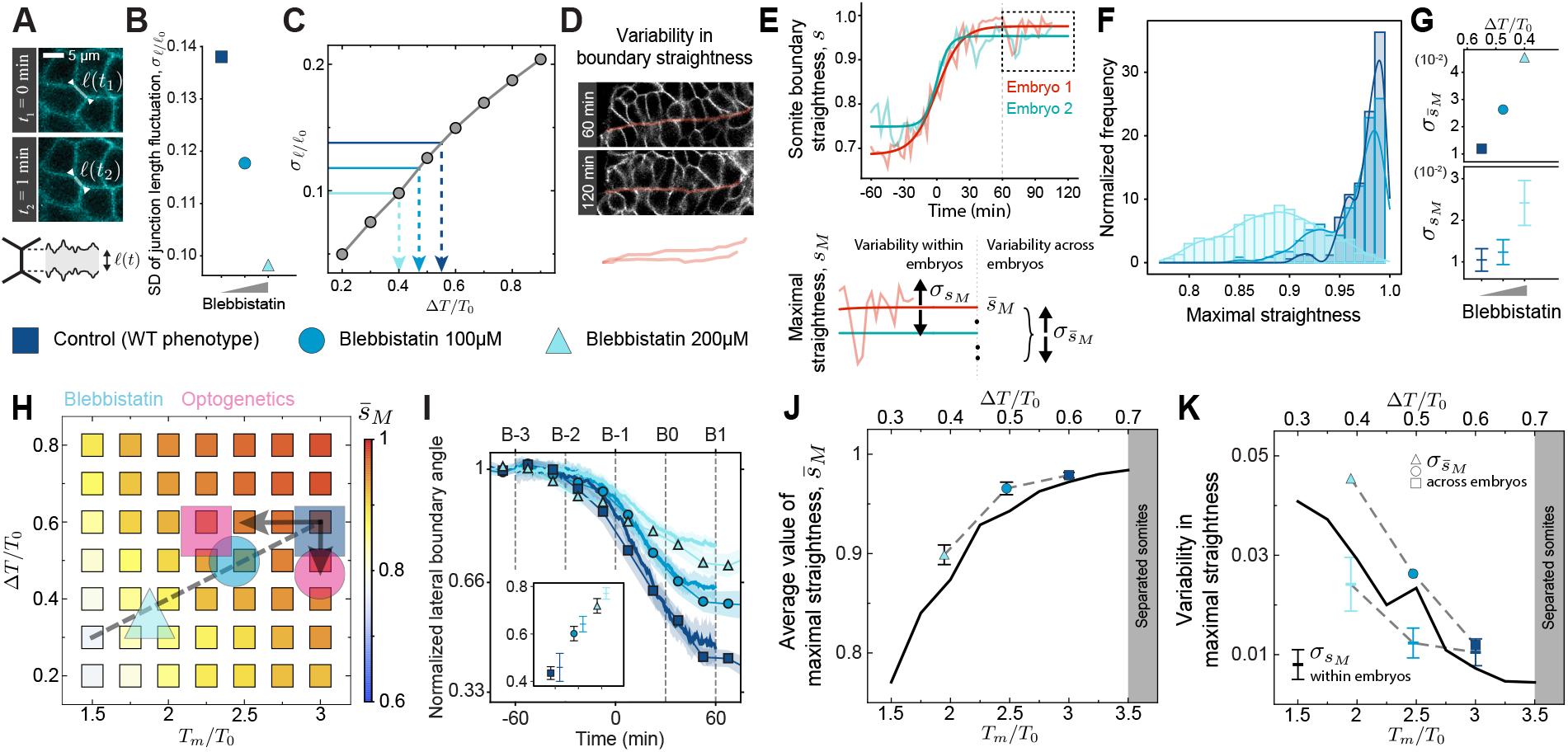
Tension fluctuations and boundary tension are optimized to define sharp somite boundaries. **A**, Confocal section of PSM cells showing junction length changes in 1 min interval. **B**, Measured standard deviation of junctional length fluctuations in control (square), blebbistatin 100µM (circle) and blebbistatin 200µM (triangle) conditions. n=365, 393, 381 junctions from 4, 4, 4 embryos, respectively. **C**, Computed standard deviation of junctional length fluctuation as a function of tension fluctuations *ΔT*/*T*_0_ (gray line and circles; for each data point, average over junctional length of 400 discrete time points from 5 independent simulations with *N* ≈ 300). Blue lines correspond experimental values measured in control embryos (dark blue) or embryos incubated in 100µM (mid blue) or 200µM (light blue) of blebbistatin. Dashed lines indicate the value of tension fluctuations for each condition obtained by comparing the computed and experimental data. **D**, Confocal section of a somite boundary showing straightness variability after saturation (t>60min). **E**, Definition of the average value of maximal straightness *s̅_M_* as well as maximal straightness variability within and across embryos, 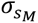 and 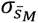 respectively. Examples of boundary straightness during somitogenesis in different embryos are shown (red, teal; both data and sigmoid fits are shown). **F**, Normalized frequency of maximal straightness *S_M_* (straightness for time≥60min) for control (dark blue), blebbistatin 100µM (mid blue) and blebbistatin 200µM (light blue). n=168, 232, 474 time points of 18, 17, 21 somites from 10, 9, 9 embryos. **G**, Measured variability of maximal straightness within and across embryos, 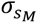 and 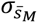 respectively, for each experimental condition. Point shapes and colors as in (**B**). n=15, 17, 20 somites from 10, 9, 9 embryos. **H**, Predicted average values of maximal straightness *s̅_M_* as a function of both the maximal boundary tension *T_M_*/*T*_0_ and the amplitude of tension fluctuations *ΔT*/*T*_0_. Square, circle and tringle show the experimental values for control embryos and increasing blebbistatin concentrations, with a gray dashed line defining its path in parameter space. Pink square and circle correspond to optogenetic perturbations of contractility at the forming boundary (square) and in the PSM (circle) (see Fig. 6F,H). **I**, Simulated (continuous line) and measured (data points and line connecting them) temporal evolution of the lateral boundary angle during straightening for the different conditions. Inset shows lateral angle evolution at B1 for experimental (shaped points) and simulated (colored bars) boundaries. **J-K**, Simulated average values of maximal straightness (**J**; continuous line) and variability of maximal straightness (**K**; continuous line) along the blebbistatin-path (gray dashed line in **H** and Fig. S8B). Measured average values of maximal straightness (**J**; shaped points connected by gray dashed lines) and measured variability of maximal straightness (**K**; 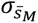 and 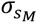) are shown. Unless otherwise stated, error bars and bands = SE.

Such tension fluctuations (mechanical noise) lead to variability in boundary straightness during somite formation (Fig. 5D-E), even after its boundary straightness saturates to maximal levels, *S_M_* (for t>60 min). To understand to origin of this variability, we decoupled the variability in maximal straightness *S_M_* arising from active mechanical noise in a given embryo (intrinsic variability), namely 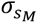, and the variability in maximal boundary straightness across embryos, 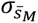 (Fig. 5E). The measured distribution of maximal boundary straightness showed a strong dependence on blebbistatin concentration (Fig. 5F; Methods), with both intrinsic variability (standard deviation, Fig. 5G, or unbounded distribution width, Fig. S10) and variability across embryos increasing with increasing levels of mechanical noise for different blebbistatin levels (Fig. 5G; Methods). Comparing quantitatively the average values of maximal boundary straightness *s̅_M_* and its total variability for different blebbistatin concentrations to the theoretical predictions showed the path of this perturbation in the parameter space spanned by *T_M_*/*T*_0_ and Δ*T*/*T*_0_ (Fig. 5H and Fig. S8B; Methods). Using these fit parameters for maximal straightness, it was possible to quantitatively explain the entire dynamics of the lateral boundary angle during somite formation for different blebbistatin concentrations (Fig. 5I; Methods). Plotting the theoretical predictions for the average value of maximal boundary straightness *s̅_M_* (Fig. 5J) and its variability (Fig. 5K) along the blebbistatin-path (Fig. 5H), together with their measured values, showed quantitative agreement between theory and experiments. While the variability across embryos 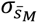 was larger than the intrinsic variability 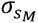 when myosin II activity was inhibited with blebbistatin (Fig. 5K), with both variability measures bounding the theoretical prediction, they became equal for wild type embryos and also equal to the theoretically predicted value, which is very close to the minimal variability possible. These results show that increasing either boundary tension or mechanical noise any further in wild type embryos does not significantly straighten the boundary or reduce its variability anymore (Fig. 5J-K). This indicates that cells at the somite boundary increase the boundary tension to just the necessary value to fully straighten the boundary, with cells in the surrounding tissue helping sharpen it by generating an optimal level of mechanical noise, thereby ensuring robust body segmentation.

To decouple the effect of boundary tension increase and active mechanical noise on boundary straightness, we perturbed RhoA activity using optogenetics, which enabled spatiotemporally controlled inhibition of myosin II activity. When illuminated, membrane bound iLID-Venus-CAAX was activated and recruited SSPB-mScarlet-DNRhoA to the cell boundary (Fig. 6A; Methods), as shown by the increased ratio of boundary to cytoplasmic signals (Fig. 6B-C). Laser ablation of cell junctions along the somite boundary in both illuminated and non-illuminated conditions showed a substantial decrease in tension upon illumination (Fig. 6D-E). In contrast, no change in junctional tension was observed in cells of the PSM or somite interior upon illumination (Fig. 6D-E), in agreement with our results using blebbistatin inhibition (Fig. 3B). However, junctional fluctuations of PSM cells or cells in the somite interior decreased following illumination (Fig. 6F), showing that mechanical noise can be modulated with optogenetics. Illuminating either the somite boundary or the interior of the somites (but not the boundary) independently, led to a decrease in boundary straightening in both cases (Fig. 6G-H). These spatially controlled perturbations allowed us to separate the effect of a reduction in boundary tension alone (boundary activation; Fig. 6G-H) or a reduction in mechanical noise in the surrounding tissue (somite interior or PSM activation; Fig. 6G-H), and trace their orthogonal paths in parameter space (Fig. 5H and Fig. S8B; magenta square and circle, respectively). Our optogenetic data indicate that both the increase in boundary tension and the active mechanical noise in the surrounding tissue are independently necessary to define sharp boundaries.

**Figure 6:**
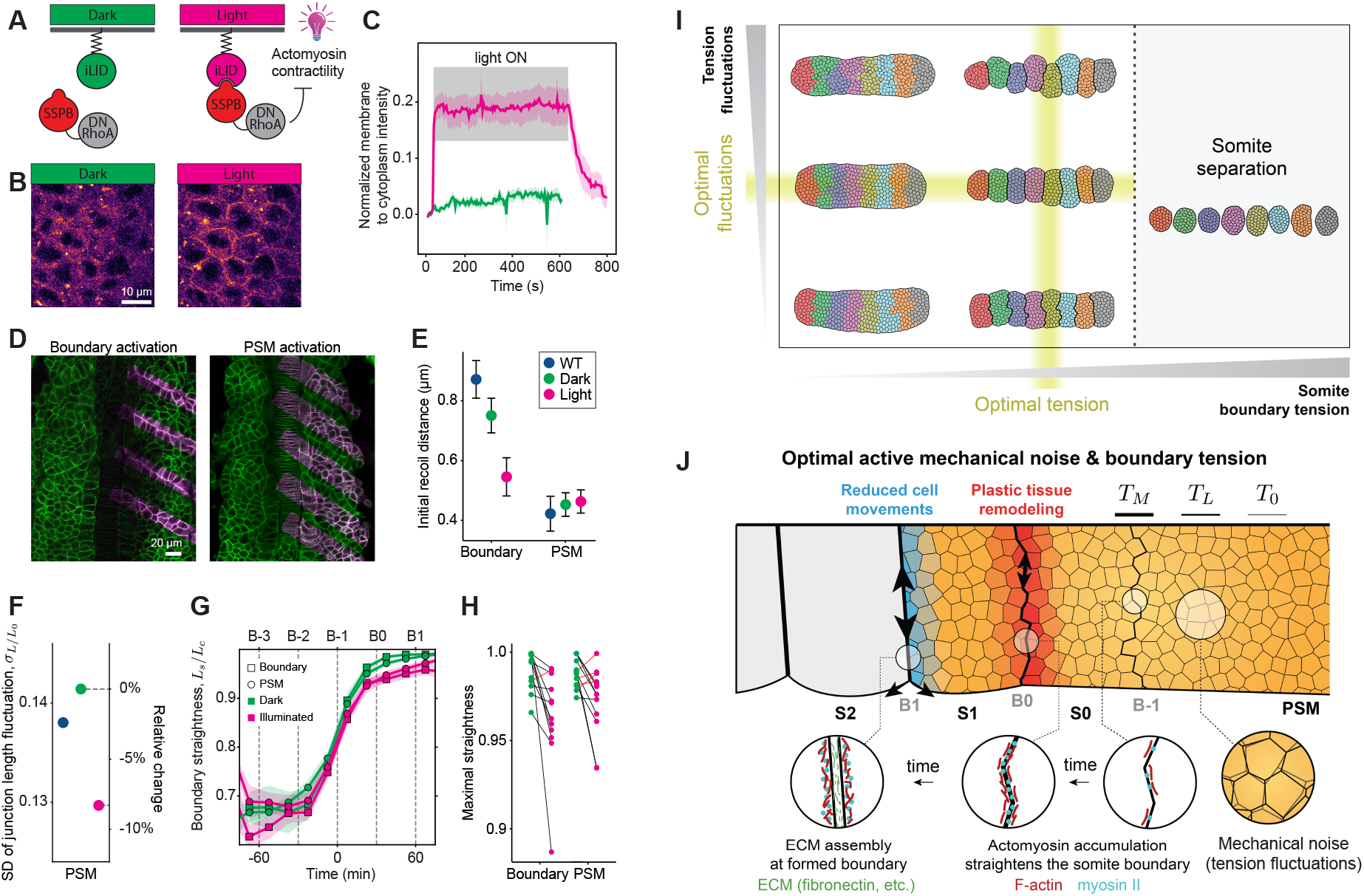
Optogenetic perturbations of somite boundary tension and mechanical noise. **A**, Schematic of iLID SSPB system. Upon blue light illumination, membrane anchored iLID gains affinity for cytoplasmic SSPB bound to DN RhoA and recruits it to the membrane where DNRhoA inhibits actomyosin contractility. **B**, SSPB-mScarlet-DNRhoA distribution in dark and illuminated conditions. **C**, Temporal evolution of normalized membrane to cytoplasmic SSPB-mScarlet-DNRhoA intensity (Methods) in dark and illuminated conditions (green and magenta curves, respectively). Gray band indicates illumination. n=6 embryos. **D**, Embryos expressing the iLID/SSPB optogenetic system. Magenta boxes show iLID activation by blue light excitation at the forming boundary or in the non-boundary mesoderm (PSM and somite interior). **E**, Initial recoil velocity after laser ablation of individual cell junctions in the somite interior or at the forming boundary for each condition: control (blue dots, from Fig. 4B), dark (green dots) and illuminated (magenta dots). n=106 & 152, 115 & 136 ablated junctions for boundary and somite interior in dark and illuminated conditions, respectively, from 21 embryos measured for both conditions. **F**, Standard deviation of junctional length fluctuations in WT embryos (blue, from Fig. 5B) and embryos expressing the iLID/SSPB optogenetic system in dark (green) and illuminated PSM (magenta) conditions. n=478, 490 junctions respectively; 5 embryos. **G-H**, Temporal evolution of boundary straightness *L_s_*/*L_c_* (**G**) and maximal straightness after straightening at B1 (**H**) at the somite boundary (squares) and in the somite interior (circles) in illuminated (magenta) and dark (green) conditions. n=12 & 13 somites; 4 embryos. Error bars = SE. **I**, Outcomes of somite formation indicating that the wildtype phenotype occurs for optimal values of boundary tension and mechanical noise (tension fluctuations) in the tissue adjacent to the boundary. **J**, Sketch detailing the main events involved in segmentation and somite formation. A progressive accumulation of F-actin and myosin II at the nascent posterior boundary of the forming somite (S0) causes an increase in boundary tension, driving its straightening, which causes a plastic remodeling of the tissue adjacent to the boundary. Once formed and straight, ECM components accumulate at the boundary and cells adjacent to the formed anterior boundary become less mobile (less rearrangements). Optimal mechanical noise (actomyosin-generated tension fluctuations) facilitate the process, enabling robust somite boundary formation. Somites are physically sectioned off the PSM by the boundary tension increase in a tissue with optimal tension fluctuations. Error bars and bands = SE.

## DISCUSSION

Altogether, our results show that the remarkable straightness of somite boundaries necessary to robustly segment the vertebrate body axis results from an optimal combination of actomyosin-generated boundary tension and mechanical noise in the tissue (Fig. 6I). The increase in tension at the somite boundary causes a transient plastic remodeling of the immediately adjacent tissue, driving boundary straightening and the physical segmentation of the body axis (Fig. 6J). Optimal mechanical noise in the tissue surrounding the somite boundary, introduced by active cell-cell contact tension fluctuations, helps robustly define maximally straight somite boundaries with minimal variability.

The notion that mechanics can help build robust structures during embryonic development was recently put forward in the context of lateral symmetry between somites^48^. Our results reveal a different role for mechanics in enabling robust body axis segmentation, in which the magnitude of active tension fluctuations in the tissue appears to be optimally tuned to enable maximal somite boundary straightening with minimal variability. Noise is typically thought to introduce defects, impairing robustness. In contrast to this notion, we find that cells actively introduce mechanical noise to facilitate the cell rearrangements necessary to sharpen the somite boundary, thereby helping the increasing boundary tension to fully straighten the boundary. It is not possible to reproduce the observed sharp boundaries by simply increasing boundary tension without active mechanical noise. This is because increasing boundary tension further would lead to the complete separation of somites (Fig. 6I). It appears that active mechanical noise in the tissue surrounding the boundary is necessary to lower the boundary tension necessary to achieve sharp boundaries without completely dissociating somites. In addition to their role in controlling tissue fluidization^39^ and cell differentiation^49^, tension fluctuations play also an important role in enabling robustness of morphogenetic processes during embryonic development.

Actomyosin accumulation at the boundary causes the initial increase in boundary tension that drives boundary straightening, as it is believed to occur in the formation of other tissue boundaries. Our measured temporal sequence of molecular assembly at the somite boundary is consistent with Eph-ephrin, actomyosin and N-cadherin interplaying to define the boundary^50–52^. While N-cadherin is not essential to form somite boundaries, its decrease at the boundary may be important to allow ECM components to assemble there, in agreement with previous observations^53^. Moreover, beyond N-cadherin and Papc, other adhesion molecules could play a role in somite formation. Our data suggest a two-step segmentation process, with actomyosin force generation driving boundary straightening and somite segmentation, followed by ECM accumulation to stabilize the somite-somite contacts^54^. A role for ECM in boundary maintenance has been previously suggested in other vertebrate tissues^25,27,28,53^, and may be essential in cases where boundary tension is used for segmentation rather than boundary stabilization.

While our simulations of the physical process of somite formation reproduce both elongated somites that maintain contact, as observed in zebrafish, as well as round separated somites, resembling those observed in trunk organoids^55^, it remains to be seen if this physical mechanism of somitogenesis is shared across vertebrate species. The observed actin distribution in somites of amniote species^22^, as well as their cellular dynamics^56^, suggest a different mechanism of segmentation from that reported here for zebrafish. Future measurements of mechanics in various species will reveal if the physical mechanism of segmentation is conserved and if the presence of active mechanical noise is also necessary to ensure robust body segmentation in other organisms.

More generally, our findings reveal how active mechanical noise, modulated to optimal levels, enables the robust formation of functional structures during embryonic development.

## RESOURCE AVAILABILITY

### Lead contact

Requests for further information and resources should be directed to and will be fulfilled by the lead contact, Otger Campàs.

### Materials availability

Newly generated reagents in this study are available from the lead contact upon request.

### Data and code availability

#### Data availability

Source data supporting these findings are available upon request.

#### Code availability

The simulations code used in this article is publicly available on GitHub at: https://github.com/campaslab/active_foam_somitogenesis

Any additional information required to reanalyze the data reported in this paper is available from the lead contact upon request.

## ACKNOWLEDGMENTS

We thank all members of the Campàs group for their comments and help, and B. Shelby, G. Stooke-Vaughan, S.-T. Yen and the UCSB Animal Research Facility for help with zebrafish care. We also thank Sean Megason (Harvard University) for kindly providing the Tg(actb2:memCherry2)^hm29^ transgenic line, C.-P. Heisenberg (IST Austria) for kindly providing the Tg(*actb2*:myl12.1-eGFP) and Tg(*actb2*:mCherry-Hsa.UTRN) transgenic lines and Tg(*actb2*:mCherry-Hsa.UTRN) transgenic lines and Scott Holley (Yale University) for kindly providing the pCS2 zf itg5a emGFP plasmid. The Tg(actb2:mem-neonGreen-neonGreen)^hm40^ line was generously provided before publication by Toru Kawanishi and Ian Swinburne in Sean Megason’s lab (Harvard University). A.B. is supported by a postdoctoral fellowship of the Alexander von Humboldt Stiftung. This work was supported by the National Institute of General Medical Sciences (R01GM135380 to EMS and OC) and the Eunice Kennedy Shriver National Institute of Child Health and Human Development of the National Institutes of Health (R01HD095797 to OC). We acknowledge support from the Center for Scientific Computing from the CNSI, MRL: an NSF MRSEC (DMR-1720256) and NSF CNS-1725797. The project was also supported by the Deutsche Forschungsgemeinschaft (DFG, German Research Foundation) under Germany’s Excellence Strategy – EXC 2068 – 390729961– Cluster of Excellence Physics of Life of TU Dresden.

## AUTHOR CONTRIBUTIONS

Conceptualization, ERS, AB and OC; Methodology, AB, ERS, SK, OC, IL and EMS; Investigation, ERS, AB and OC; Formal analysis, AB, ERS, SK, BG, MP and RW; Software, SK; Writing – original draft, ERS, AB and OC; Visualization, ERS, AB and OC; Funding acquisition, AB, EMS and OC; Supervision, OC.

## SUPPLEMENTAL INFORMATION

Figures S1–S10; Supplementary Movies 1–5; Table S1 (simulation parameters).

## STAR★METHODS

### KEY RESOURCES TABLE

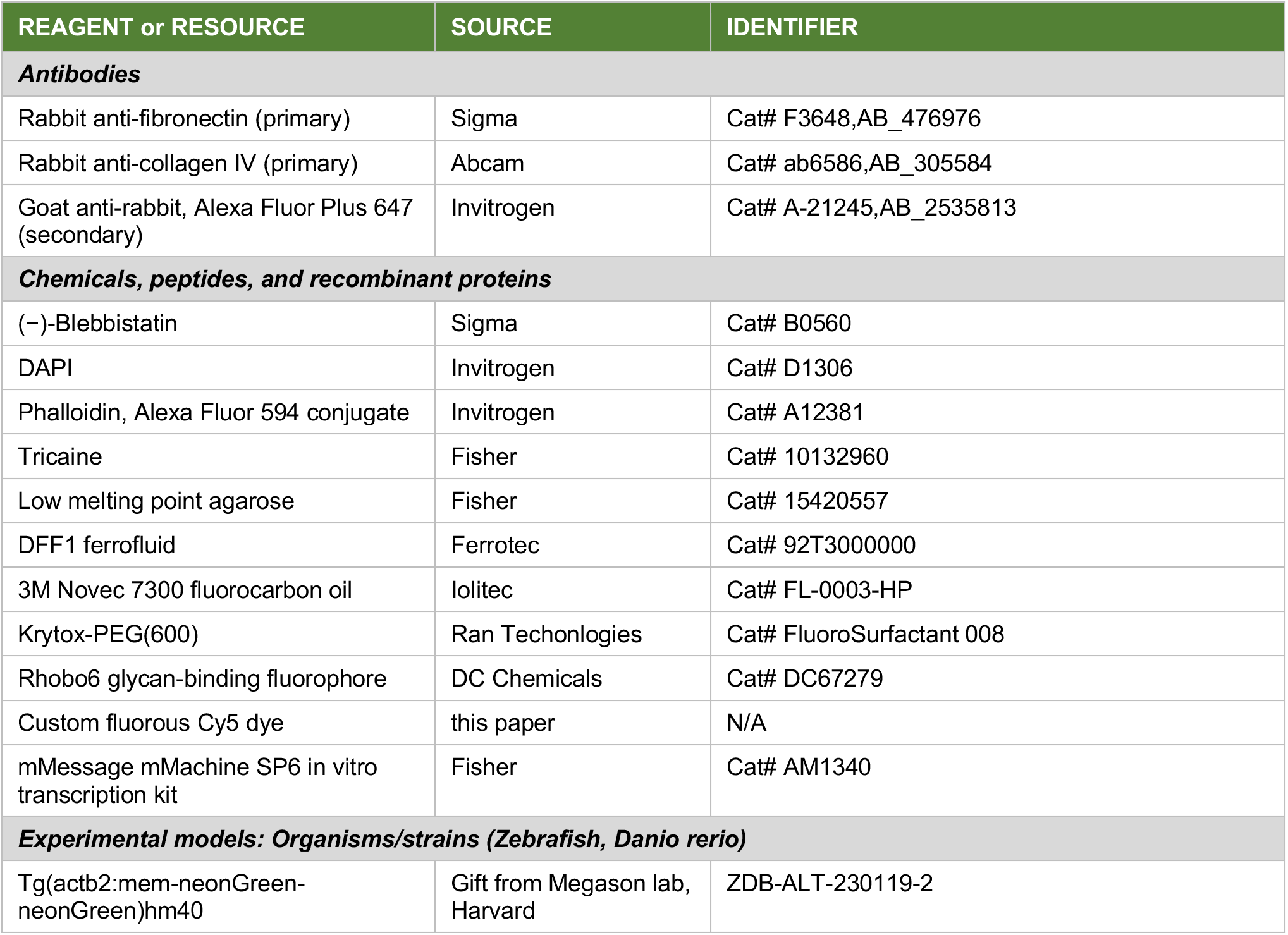
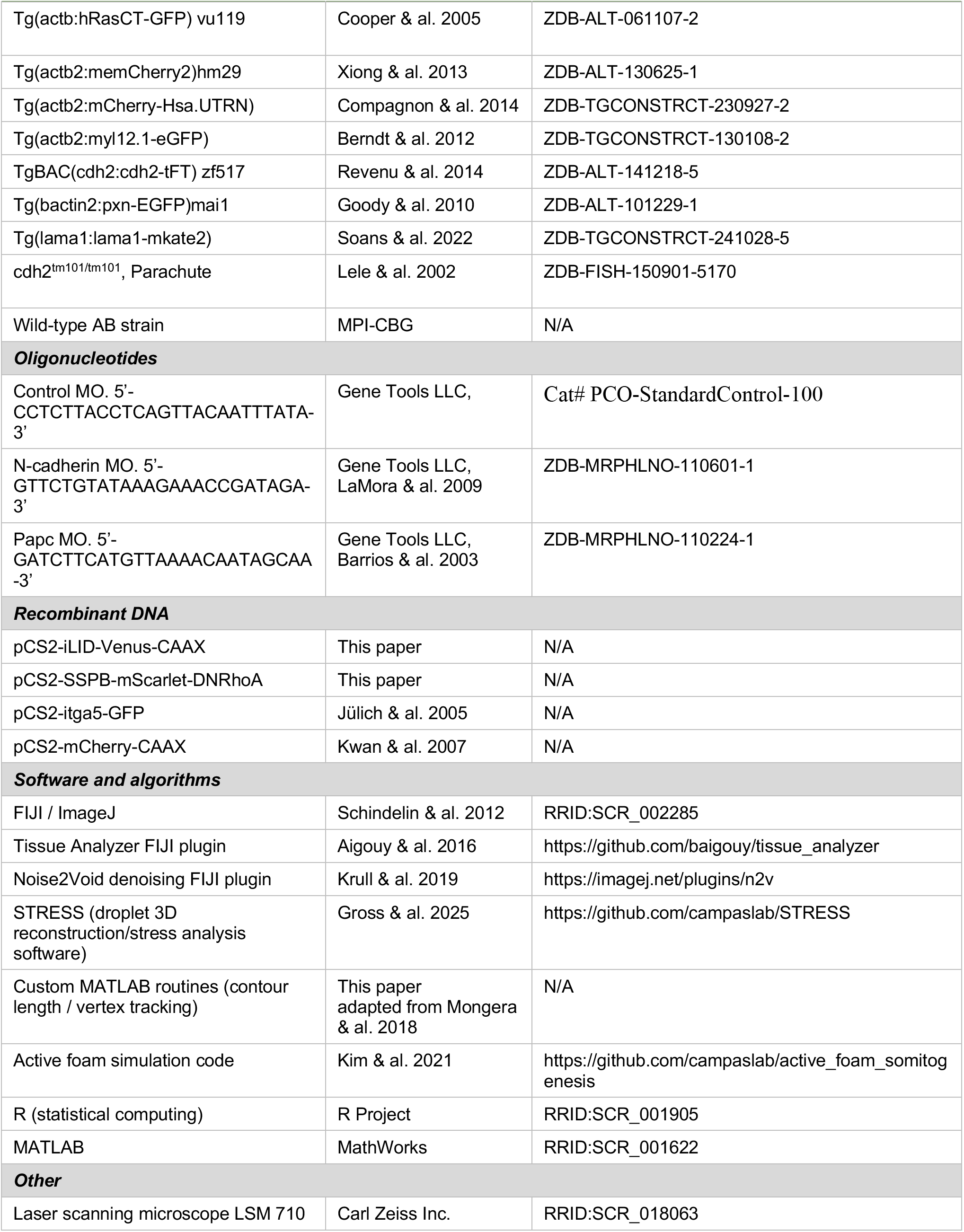
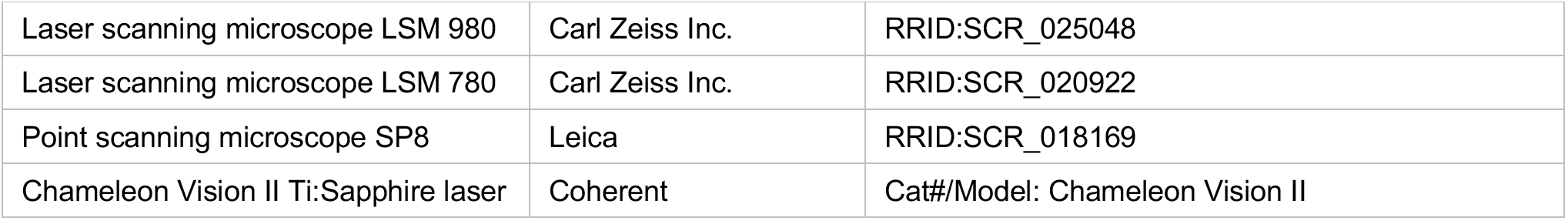

## EXPERIMENTAL MODEL AND STUDY PARTICIPANT DETAILS

### Zebrafish husbandry and transgenic lines

Zebrafish (*Danio rerio*) were maintained as previously described^57^. Animals were raised and experiments were performed following all ethical regulations and according to protocols approved by the Institutional Animal Care and Use Committee (IACUC) at the University of California, Santa Barbara, and by the European Union (EU) directive 2010/63/EU as well as the German Animal Welfare act. Transgenic lines *Tg(actb2:mem-neonGreen-neonGreen)^hm40^*, *Tg(actb:hRasCT-GFP)*^58^ or *Tg(actb2:memCherry2)^hm29^*^59^ were used to visualize cell membranes. *Tg(actb2:mCherry-Hsa.UTRN)* ^60^, *Tg(actb2:myl12.1-eGFP)* ^61^, *TgBAC(cdh2*:*cdh2*-*tFT)* ^62^, *Tg(bactin2:pxn-EGFP)^mai1^*^63^ and *Tg(lama1:lama1-mkate2)* ^64^ were used to visualize F-actin and myosin II, N-cadherin, paxillin and laminin α1, respectively. Mutant line cdh2^tm101/tm101^ (also called Pac^65^) was used to observe somite boundary formation in an N-cadherin KO line.

## METHOD DETAILS

### Morpholino and mRNA injection

Embryos obtained using an outcross of *Tg(actb2:myl12.1-eGFP)* and Tg(actb2:memCherry2)^hm29^ were injected at the 1-cell stage with 0,4 pmol of standard control morpholino (5’-CCTCTTACCTCAGTTACAATTTATA-3’) or N-cadherin targeting morpholino (5’-GTTCTGTATAAAGAAACCGATAGA-3’) ^66^ and/or 0.1 pmol of Papc targeting morpholino^45^ (5’-GATCTTCATGTTAAAACAATAGCAA-3’) (Gene Tool LLC, Philomath). To visualize integrin α5, we injected mRNA encoding for itga5-GFP (150pg) ^67^ and mCherry-CAAX (100pg) in AB embryos at one cell stage. To express the SSPB/iLID optogenetic system, we injected mRNA encoding for iLID-Venus-CAAX (250pg) and SSPB-mScarlet-DNRhoA (25pg) in AB embryos at one cell stage. mRNAs were synthetized using the mMessage mMachine SP6 in vitro transcription kit (Ambion).

### Fibronectin and collagen IV labelling in fixed embryos

Embryos of the *Tg(actb2:myl12.1-eGFP)* line for immunostaining were manually dechorionated at 12 somites stage and fixed in 4% paraformaldehyde overnight at 4°C and then rinsed in phosphate-buffered saline with 1% Triton X-100 (PBT). Embryos were permeabilized by incubating in PBT with 0.06% trypsin for 4 min and rinsing in PBT. Embryos were then incubated overnight at 4°C with primary antibody against fibronectin (Sigma) or collagen IV (Abcam) then rinsed in PBT. Finally, embryos were incubated overnight at 4°C in secondary antibodies goat anti rabbit conjugated to Alexa fluor plus 647 (Invitrogen), 1:1,000 DAPI and Phalloidin conjugated to Alexa fluor 594 (Invitrogen) diluted in PBT and then rinsed in PBT.

### Drug treatment

Embryos used in drug treatment experiments were dechorionated around the 8 somites stage and incubated in a solution (-)-Blebbistatin (Sigma-Aldrich) (100µM or 200µM), or DMSO (1%) in E3 for at least one hour at 25°C. After being mounted in agarose for imaging, embryos were covered by the incubation solution.

### Imaging

Embryos were mounted in 1% low melting point agarose (E3 media containing 0.01% Tricaine) at approximately the 10 somite stage and imaged at 25°C. Embryos were imaged using the following combination of microscopes and objectives: Laser scanning microscope (LSM) 710 or LSM 980 (Carl Zeiss Inc.) with a 40x water immersion objective (LD C-Apochromat 1.1 W, Carl Zeiss), LSM 780 with a 40x water immersion objective (LD C-Apochromat 1.2 W, Carl Zeiss), or point scanning SP8 microscope (Leica) with a 40x water immersion objective (1.1 HC PL IRAPO, Leica). Images were acquired at 15 second intervals for *in situ* droplet actuation experiments and at 1 to 3 minute intervals for timelapse imaging of somite formation, depending on the experiment. All imaging of embryos was done using a 40x water immersion objective (LD C-Apochromat 1.1 W, Carl Zeiss). Imaging of protein fluorescence intensity was done with spatial resolution ranging from 0.25-0.69 µm. Volumetric images of droplets were done at 0.35µm spatial resolution, and 4.0 µm z-steps. After timelapse imaging, embryos were removed from agarose and imaged laterally to determine the position of the droplet along the AP axis. All stress measurements were performed between the 14 and 16 somites stages. For measuring junction length fluctuations, embryos were imaged for 5 minutes every 5 seconds with a 0.11µm spatial resolution using the Airyscan detector of a LSM 980 (Carl Zeiss).

### Quantification and statistical analysis

The statistical analyses were performed using R and MATLAB. Statistical details of experiments are reported in the figures and figure legends. Sample size is reported in the figure legends. No statistical test was used to determine sample size. The biological replicate is defined as the number of cells, somites and/or independent embryos, as stated in the figure legends. No inclusion/exclusion or randomization criteria were used and all analyzed samples are included. Unless differently stated in the figure legends, the graphs show mean ± SEM and the error bars are calculated and shown based on the number of cells, somites or embryos, as indicated.

### Quantification of somite boundary morphology

Forming somite boundaries were identified and annotated manually using sequences of confocal sections of Tg(*actb2*:mem-neonGreen-neonGreen)^hm40^, Tg(actb:hRasCT-GFP) or Tg(*actb2*:memCherry2)^hm29^ embryos. First, we identified the location of the mature somite boundary in the final frame of the timelapse sequence. Next, we identified the boundary in the previous frame in the sequence, using the most recently analyzed frame as a reference. At each frame, boundary annotations were recorded as ordered sets of coordinates. The boundary contour length *L_c_* was computed by summing the length of line segments connecting the coordinates along the contour. The straight end-to-end distance of the somite boundary, *L_s_*, was computed as the distance between the first and last coordinates of the contour. Somite-somite boundaries were annotated at 1 minute or 3 minute increments depending on the experiment, whereas lateral boundaries were traced at 15 minutes intervals. Adaxial cell boundaries were not included in boundary annotations. Lateral boundary angle was identified after establishing the trace of somite boundary as the angle formed by the two most distal cells, on each side of the boundary. Lateral boundary angle was manually annotated and measured at 3 minutes increments using the Angle tool of FIJI^68^.

### Temporal registration of different embryos and staging of somite formation

Data sets collected from different embryos were temporally aligned with each other using the temporal evolution of the straightness measure, namely *m_s_*(*t*) *≡ L_s_/L_c_*. For each somite boundary in each experiment, the measured straightness was fit to the function *m_s_*(*t*) = *A/*(1 + *e*^(*t−B*)*×C*^) + *D*, where *A* specifies the amplitude, *B* specifies time of half-time between initial and final states, *C* specifies the ramping rate, and *D* specifies the baseline offset. For each embryo, we associated the half-time *B* with the time of stage B-1 of somite boundary formation (Fig. 1B), as the straightening corresponds to the nascent posterior boundary of somite S0^69^. Experiments in different embryos were then temporally aligned by registering their half-times and shifting them all to t=0. Therefore, t=0 corresponds to the stage B-1 of somite boundary formation. The stages S(N) of somite formation (S0 referring to the currently forming somite, S1 referring to the most recently formed somite, and so on) and their posterior (caudal) and anterior boundaries, namely B(N-1) and B(N) respectively, were denoted following established nomenclature^69^ (Fig. 1B). Boundaries at stages B(N-1) (N = −2, −1, 1, 2,…) occurred at times *t* = *N T* relative to stage B-1 (*t* = 0 min), with *T* = 30.0 *±* 2.7 min being the period of somite formation in our experiments (wild type zebrafish embryos at 25 °C^70^).

### Quantification of protein levels at somite-somite and lateral boundaries

The following outcrosses were used to simultaneously label membrane and a protein of interest. Tg(*actb2*:mCherry-Hsa.UTRN) and Tg(*actb2*:mem-neonGreen-neonGreen)^hm40^ to visualize F-actin. Tg(*actb2*:myl12.1-eGFP) and Tg(*actb2*:memCherry2)^hm29^ to visualize myosin II. Tg(bactin2:pxn-EGFP)^mai1^ and Tg(*actb2*:memCherry2)^hm29^ to visualize paxillin. Tg(lama1:lama1-mkate2) and Tg(actb:hRasCT-GFP) to visualize laminin α1. Quantification of the N-cadherin signal at the somite boundary was obtained using an incross of TgBAC(*cdh2*:*cdh2*-*tFT*) to visualise N-cadherin which is accumulated at cell-cell junctions, directly allowing cell contour visualization. Integrin α5 signal at the somite boundary was quantified using injection of mRNA encoding for itg5a-GFP and membrane bound mCherry in AB embryos to visualize integrin α5 and membranes simultaneously. In order to non-specifically label ECM proteins, 0.4pmol of the glycan binding fluorophore Rhobo6^46^ was injected in the extracellular space of *Tg(actb:hRasCT-GFP)* embryos at the 8 somites stage.

Intensities of the different fluorescent proteins and fluorophores along the boundary contour were computed from boundary coordinates (see above) and fluorescent images sequences using custom Matlab scripts. Traced boundary coordinates were converted into a Polyline ROI object, which was then converted into a binary array using the function createMask(). This binary array was then dilated with a disk shaped structuring element using the function imdilate(), resulting in boundary masks approximately 3 µm in width. These masks were used at each frame to compute the average signal intensity along the boundary at each timeframe. We determined if considerable photobleaching occurred in a given timelapse by quantifying the temporal evolution of the average signal intensity within control rectangular regions (of area ≃3200 µm^2^) in the PSM, where the average intensity is not supposed to change in time. If significant bleaching was observed for a given sequence, a single exponential photobleaching curve was determined by nonlinear regression and a time dependent bleach correction factor was applied to measurements from that sequence. After photobleaching correction (if required), signal intensities at the somite boundary were normalized by the average intensity in the somite boundary before somite formation starts, namely within the [-60,-30] min time interval (*t* = 0 min, corresponding to B-1).

### Quantification of fibronectin and collagen IV fluorescence signal at somite-somite and lateral boundaries

Quantification of the fibronectin signal at the somite boundary was obtained by performing immunostaining against fibronectin and collagen IV in *Tg(actb2:myl12.1-eGFP)* embryos fixed at the 12 somites stage. Co-staining for actin allowed visualizing cell contour while labelled myosin II GFP allowed to locate the forming somite boundary as a line of accumulated myosin signal. To assign a developmental time to the different somites, we measured the straightness of the forming somite boundary (somite B-1) and compared it to the straightening curve measured on control, living embryos. In particular, we assigned to the forming boundary in the fixed sample the developmental time by matching its straightness value to the reference straightening curve. Then, we considered each side of every embryo as a series of somites. Every other somite boundary on the same series is staged 30 min older than the preceding. By aggregating the different series of somite boundaries, we could establish a curve of straightening over time very close to those measured in live experiments. Finally, we measured the intensity of fibronectin for each somite boundary and, for each series, somite intensity is normalized between 0 and 1, using the following formula: *I_s_*_,*norm*_ = (*I_s_* − *I_min_*)⁄(*I_max_* − *I_min_*). With *I_s_* the measured fibronectin or collagen IV intensity of somite s, *I_s_*_,*norm*_ the normalised fibronectin intensity of somite s, and *I_min_* and *I_max_* the minimal and maximal values of fibronectin or collagen IV intensity measured on this series of somite boundaries, respectively.

### Kymograph visualization of F-actin and myosin II signals at the somite boundary

For each column corresponding to a time point in the sequence, the fluorescent actin (or myosin) signal was measured along the somite boundary by averaging intensities within a 3 *µ*m diameter circular mask at an increment of 3 *µ*m along the path specified by the annotated boundary. The kymograph shows the traced intensities along the boundary contour for each time frame (frame rate = 3 min).

### Generation and injection of ferrofluid droplets

Ferrofluid droplets were prepared as previously described^34,35^. Briefly, DFF1 ferrofluid (Ferrotec) was diluted in filtered 3M Novec 7300 fluorocarbon oil. To prevent non-specific adhesion between cells and droplets, a fluorinated Krytox-PEG(600) surfactant (008-fluorosurfactant, RAN Biotechnologies^71^) was diluted in the ferrofluid at a 2.5% (w/w) concentration. A custom-made fluorous Cy5 dye was used to visualize the droplet (see below). The ferrofluid was calibrated before each experiment as previously described^35^, so that the applied magnetic stresses are known. Once prepared and calibrated, the ferrofluid was injected into the lateral mesodermal progenitor zone between 4 and 6 somites stage to form droplets of approximately 30 µm in diameter, as previously described^34,35^. Imaging of droplets started at least 1.5 hours after the injection to let the tissue fully recover from it. Embryos were incubated for 2 hours at 25*^◦^*C following injection to allow for tissue recovery, as previously described^35^.

### Magnetic actuation of ferrofluid microdroplets

Actuation of ferrofluid droplets was performed as previously described^35^. Briefly, ferrofluid droplets were actuated by a uniform and constant magnetic field to deform the droplet and apply stresses in the surrounding tissue. The magnetic field was applied for 20 minutes and subsequently turned off. The droplet interfacial tension *γ* was measured in each experiment *in situ* and *in vivo* from these actuation experiments, as previously described^34,35^.

### Generation of fluorinated cyanine dye for droplet imaging

To visualize ferrofluid droplets and perform 3D stress measurements *in vivo* and *in situ*, we employed a custom-synthesized fluorous Cy5 dye. While fluorous Rhodamine dyes have been previously used to visualize droplets^34,35^, longer wavelength dyes are preferred for full 3D reconstructions of droplets. The fluorous Cy5 was synthesized using a similar protocol as previously described^72^. Briefly, a branched fluorous ketone was reacted with phenyl hydrazine to produce a fluorous indole. A fluorous methylene indolene was generated by *N*-alkylation with a perfluoroalkyl iodide. The heterocycle was condensed onto malonaldehyde bis(phenylimine) monohydrochloride in the presence of a pyridine base in acetic anhydride to generate the fluorous Cy5. The dye was isolated by column chromatography in 9% yield. The synthesized fluorous Cy5 dye was then diluted in the ferrofluid at a final concentration of 25 *µ*M. We have previously show that the dye is biocompatible and enables robust 3D measurements of mechanical stresses^72^.

### Measurements of cell-scale and tissue-scale stresses

Stresses were quantified from the deformations of droplets inserted in the tissues, as previously described^34,37,73^. Briefly, droplets were imaged in 3D using confocal microscopy and their shape was reconstructed in 3D using automated software^36,37^. Using the measured value of the interfacial tension for each droplet (see above) and the time evolution of the droplet geometry, we measured the time evolution of cell- and tissue-scale stresses, as detailed in reference^37^. Tissue-scale stresses were obtained from the ellipsoidal deformation mode of the droplet, whereas cell-scale stresses were linked to higher-order modes after removing the ellipsoidal mode (Fig. S7A). Reported amplitudes of cell-scale stresses were obtained using the value *α* = 0.05 in the analysis software to remove the smallest and largest 5% values of stresses, as these extreme values are prone to noise.

### Measurement of stresses at somite-somite boundaries

The magnitude of stress anisotropy generated at the somite boundary was quantified by comparing the stresses along the somite boundary to the stresses in the direction perpendicular to the somite boundary. To do so, we measured the stresses in different surface regions on the droplet. At each time point, we defined a belt region as the set of droplet surface coordinates located within 5 µm of the somite boundary plane, which was identified by hand at each timepoint. We also defined anterior and posterior caps as the two 10 µm diameter regions on the droplet most distant from the somite-somite plane.

Let *H_B_*, *H_A_*, and *H_P_* refer to the sets of mean curvature measurements within the belted region, the anterior cap, and the posterior cap, respectively. The boundary stress, *σ_B_*, reads

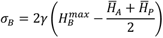

where *γ* is the droplet interfacial tension, 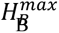 is the maximal mean curvature value within the boundary belt, and *H̅_A_* and *H̅_P_* are the average mean curvatures in the anterior and posterior caps, respectively.

### Quantification of T1 transition (neighbor exchange) rates and orientation

To measure the spatiotemporal distribution in T1 transitions during somite formation, embryos featuring a membrane label were imaged at 1 minute intervals for at least 2 hours, as described above. Temporal alignment relative to B-1 was determined as described above. Each sequence was first divided into 10 minute increments, and then a quadrilateral grid was defined on the somite at that interval, as shown in Fig. 3A,B. The grid divides a forming somite into 9 segments along the ML direction and 5 segments along the AP direction. Moreover, the grid is extended further two rows anteriorly and posteriorly. All observed T1 transitions were recorded within the grid for each sequence, from S-2 to S2 and classified in 10 minute increments. The grid was used to define three regions within a forming somite: anterior boundary, somite interior, and posterior boundary. The anterior boundary region is the anterior-most row located within the somite, the posterior boundary region is the posterior-most row located inside the somite, and the somite interior region corresponds to the inner three rows between the anterior and posterior regions. T1 rates were computed within each region for each sequence. In total 14 somite formation sequences from 7 unique embryos were used. Measurements from all sequences were used to compute averages and standard errors.

The orientation of each T1 transition was measured as the angle formed by the junction to disappear the time before rearrangement and the somite boundary defined as the border of the grid. All these measurements were pooled to show the distribution of in plane T1 transition. Rearrangements where cell disappears from the plan are counted as out of plan T1 transition. The percentage of out of plan rearrangements in the total amount of T1 transitions was measured for each embryos individually and used for computing averages and standard errors.

### Laser ablation

Laser ablation experiments were performed using a LSM 780 (Carl Zeiss Inc.) equipped with a 40x water immersion objective (LD C-Apochromat 1.2 W, Carl Zeiss) and a Chameleon Vision II tuneable pulsed near-infra-red laser (Coherent). Embryos were obtained using an incross of Tg(actb2:hRasCT-GFP) to visualize the membrane.

For ablations of cell-cell junctions, timelapses were acquired at the rate of one frame every second. Cell-cell junctions roughly perpendicular to the body axis junction in the PSM or in formed somite boundaries were selected for ablation. A rectangular ROI of 1.2 by 4.6 µm, long axis parallel to the body axis, was drawn at the middle of the junction. After 2 imaging frames, laser treatment using the pulsed laser set at 800nm & 400 mW output power was applied on the ROI, followed by 5 imaging frames. Movies of ablated cell-cell junction were analysed using FIJI. They were first denoised using the N2V plugin^74^. A line was then drawn parallel and close to the ablated junction and fluorescence profile along this line the frames before and after ablation were saved. Data were then processed in R. Vertices at each side of the ablated junction were automatically identified as peak of fluorescence intensity and the distance between these two peaks just before and after ablation were measured. Recoil was then computed as the difference between the distance after and the distance before ablation.

For 3D tissue laser ablation, 3D timelapse was acquired at a rate of one Z-stack every minute for at least 15 min prior to ablation. A square ROI of 8 by 8 µm was then drawn at the middle of the B0 junction on one side of the embryo. Starting from the bottom, ventral part of somitic mesoderm and up to the dorsal part, a laser treatment was applied on the ROI every 5 to 10µm to pierce a 3D hole through the forming somite-somite boundary, as described in^75,76^. After ablation, acquisition of the 3D timelapse was resumed for at least 45 min. The use of non-linear optics provided sufficient axial resolution not to affect the yolk cell underneath, nor the tissues above. The wound created by treatment healed within an hour and development of treated embryo was then undistinguishable from untreated ones. Measurement of the lateral boundary angle has been performed as described above.

### Normalized frequency (distribution) of cell–cell contact length fluctuations

To determine the characteristic timescale of changes in cell–cell contact lengths, we acquired a single confocal section of embryos injected with mCherry-CAAX mRNA for 30 min with time resolution of 5s using the AiryScan detector of the LSM 980 microscope (Carl Zeiss Inc.). We detected the location of cell vertices and cell–cell contact lengths in the images using the FIJI plugin Tissue Analyzer^77^. For each embryo, we segmented a ROI in the PSM for which cell-cell contacts were trackable over a 300 s time period. We then used published MATLAB (MathWorks) routine^34^ to obtain the contour length of cell–cell contacts and the (x, y) positions of the vertices. We selected all cell–cell contacts that persist for at least 100 s, and obtained their time-dependent normalized lengths *l*/*l*_0_, where *l* is the cell–cell contact length at a given time point and *l*_0_ is the average length of each individual cell–cell contact over time. We then combined all the values of *l*/*l*_0_ of all times and cell–cell contacts into a single, normalized frequency distribution, which peaks around 1, as described in^34^. The magnitude of tension fluctuations was obtained from the standard deviation of this distribution.

### Active foam simulation of somite formation

To model somite formation, we adapted the theoretical framework of active foam simulations^39^. For initial configurations, a rectangular slab of cells that consists of eight prospective somites is generated with open boundary conditions. Each somite contains approximately 40 cells, 8 rows of cells in the ML direction and 5 rows of cells in the AP direction. Initial somite boundary is specified by assigning a somite ID to individual cells. To match experimentally observed initial somite boundary straightness, cell ID at the somite boundary is randomly swapped until the boundary straightness matches with the experimental value at stage B-3. As this tissue region, the anterior PSM, is close to confluence, we simulated the system at confluence throughout somite formation.

The equations governing the dynamics of the system are the same as in reference^39^ and were integrated using the Euler-Maruyama method. The ramp up of heterotypic tension is implemented as a temporal increase of the target (fixed point) tension for heterotypic cell-cell contacts. The heterotypic tension ramp up was applied sequentially to somite boundaries and delayed by 30 minutes between them to match the experimentally observed time of somite formation. Throughout the simulations, T1 transitions were applied if a junctional length became shorter than a critical length, as previously described^39^. The specific parameters used in the simulations, as well as their values, are indicated in Supplementary Table 1.

### Optogenetics constructs and illumination patterns

To control cell contractility, we used the iLID/SSPB system^78^ coupled to a dominant negative from of RhoA^79^. A first construct, iLID-Venus-CAAX was obtained from Addgene^78^ and cloned into the pCS2+ vector. A second construct, SSPB-mScarlet-DNRhoA was synthetized directly in pCS2+ (Twist Bioscience) using as a template the existing construct SSPB-mCarlet-MYPT^80^ and DNRhoA^79^ (RhoA T19N & C189Y). Upon blue light illumination (445 or 458nm), iLID affinity for SSPB increase by forty to fifty folds^78^, leading to spatiotemporally restricted recruitment of SSPB. To measure the kinetics of SSPB recruitment upon illumination, we imaged SSPB distribution every 5 seconds with and without blue light activation. Venus was excited using a 514nm laser line to avoid iLID activation. We segmented membranes as described above and quantified SSPB intensity on a band 1µm wide following the membranes (membrane signal, *I_m_*), as well as two bands 1µm wide and 0.5µm away from the membranes on each side (peri-membrane signal, *I_p_*). We measured the normalized membrane to cytoplasmic SSPB signal ratio, *R_I_*, defined as *R_I_*(*t*) = (*I_m_*(*t*)/*I_m_*(*t*)) − (*I_m_*(*t*_0_)/*I_m_*(*t*_0_))/(*I_m_*(*t*_0_)/*I_m_*(*t*_0_)). We observed that iLID activation and deactivation time constants are of 8.7s and 84.7s, respectively, consistent with previous reports^78^.

We measured the effect of the optogenetic during somitogenesis as described above with the following modifications. For junction length fluctuation, WT embryos injected with the optogenetic system were imaged during 10 min using the Venus channel as a membrane marker and mScarlet to check for SSPB recruitment. In the “Blue light ON” condition, the field of view was also scanned with a 445nm laser, leading to iLID activation and SSPB recruitment. We then kept the last five minutes of each movie and quantified junction length fluctuation as described earlier.

For junctional tension, cell-cell junctions in the PSM or at the forming somite boundary in WT embryos injected with the optogenetic system were ablated. In the “Blue light ON” condition, the field of view is scanned every 5 second and during 2 minutes with a 458nm laser prior to the ablation.

For boundary straightening, 3D stacks of the PSM of WT embryos injected with the optogenetic system were acquired every two minutes. After each acquisition, iLID was activated in 3D through illumination with 445nm laser in ROIs located either at the forming somite boundaries, or in the PSM and somitic mesoderm (Fig. 5L). Position of the ROIs was adjusted every 20 minutes to account for global tissue flows.

### Quantification of maximal straightness variability

In order to quantify the different sources of variability in maximal straightness (i.e. straightness after saturation, t ≥ 60min), we defined two measures: the variability within (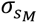) and across (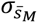) embryos (Fig. 5E, G). 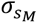 corresponds to the standard deviation of maximal straightness (i.e., value of straightness for t>60min) within embryos (intrinsic variability; Fig. 5E). This is measured for a single somite in a single embryo, and then the obtained values of the standard deviation are averaged. 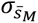 is defined as the standard deviation of the average values of maximal straightness across embryos (Fig. 5E). To show the distribution of maximal straightness, we pooled all the measurements of straightness after saturation and plotted it in a histogram grouped by experimental conditions (Fig. 5F).

### Quantification of unbounded straightness variability

Since the boundary straightness is bounded at 1, its variation (e.g., standard deviation) could be affected by the presence of this bound. In order to get an estimation of the unbounded variability of boundary straightness, we measured the width of its distribution in its unbounded side, which we call the unbounded distribution width (Fig. S10). The unbounded distribution width is the width of the distribution covering 45% of the distribution below the average (the unbounded side of the distribution). We removed 5% of the smallest values because these are prone to noise. This measure of variability reliably represents the dispersion of boundary straightness even when the average boundary straightness close to 1.

## REFERENCES

1. Collinet, C., and Lecuit, T. (2021). Programmed and self-organized flow of information during morphogenesis. Nat Rev Mol Cell Bio 22, 245–265. 10.1038/s41580-020-00318-6.

2. Boutillon, A., Banavar, S.P., and Campàs, O. (2024). Conserved physical mechanisms of cell and tissue elongation. Development 151. 10.1242/dev.202687.

3. Gavish, A., Shwartz, A., Weizman, A., Schejter, E., Shilo, B.-Z., and Barkai, N. (2016). Periodic patterning of the Drosophila eye is stabilized by the diffusible activator Scabrous. Nat. Commun. 7, 10461. 10.1038/ncomms10461.

4. Umulis, D., O’Connor, M.B., and Othmer, H.G. (2008). Robustness of Embryonic Spatial Patterning in Drosophila melanogaster. Curr. Top. Dev. Biol. 81, 65–111. 10.1016/s0070-2153(07)81002-7.

5. Giammona, J., and Campàs, O. (2021). Physical constraints on early blastomere packings. Plos Comput Biol 17, e1007994. 10.1371/journal.pcbi.1007994.

6. Yamamoto, K., and Kimura, A. (2017). An asymmetric attraction model for the diversity and robustness of cell arrangement in nematodes. Development 144, 4437–4449. 10.1242/dev.154609.

7. Hubaud, A., and Pourquié, O. (2014). Signalling dynamics in vertebrate segmentation. Nature reviews Molecular cell biology 15, 709–721. 10.1038/nrm3891.

8. Bénazéraf, B., and Pourquié, O. (2013). Formation and Segmentation of the Vertebrate Body Axis. Annu Rev Cell Dev Biol 29, 1–26. 10.1146/annurev-cellbio-101011-155703.

9. Dequéant, M.-L., and Pourquié, O. (2008). Segmental patterning of the vertebrate embryonic axis. Nat Rev Genet 9, 370–382. 10.1038/nrg2320.

10. Giampietro, P.F., Raggio, C.L., Blank, R.D., McCarty, C., Broeckel, U., and Pickart, M.A. (2013). Clinical, Genetic and Environmental Factors Associated with Congenital Vertebral Malformations. Mol. Syndr. 4, 94–105. 10.1159/000345329.

11. Eckalbar, W.L., Fisher, R.E., Rawls, A., and Kusumi, K. (2012). Scoliosis and segmentation defects of the vertebrae. Wiley Interdiscip. Rev.: Dev. Biol. 1, 401–423. 10.1002/wdev.34.

12. Pourquie, O. (2003). The Segmentation Clock: Converting Embryonic Time into Spatial Pattern. Science 301, 328–330. 10.1126/science.1085887.

13. Jiang, Y.-J., Aerne, B.L., Smithers, L., Haddon, C., Ish-Horowicz, D., and Lewis, J. (2000). Notch signalling and the synchronization of the somite segmentation clock. Nature 408, 475–479. 10.1038/35044091.

14. Palmeirim, I., Henrique, D., Ish-Horowicz, D., and Pourquié, O. (1997). Avian hairy Gene Expression Identifies a Molecular Clock Linked to Vertebrate Segmentation and Somitogenesis. Cell 91, 639–648. 10.1016/s0092-8674(00)80451-1.

15. Venzin, O.F., and Oates, A.C. (2020). What are you synching about? Emerging complexity of Notch signaling in the segmentation clock. Dev. Biol. 460, 40–54. 10.1016/j.ydbio.2019.06.024.

16. Oates, A.C., Morelli, L.G., and Ares, S. (2012). Patterning embryos with oscillations: structure, function and dynamics of the vertebrate segmentation clock. Development 139, 625–639. 10.1242/dev.063735.

17. Mara, A., and Holley, S.A. (2007). Oscillators and the emergence of tissue organization during zebrafish somitogenesis. Trends Cell Biol. 17, 593–599. 10.1016/j.tcb.2007.09.005.

18. Hubaud, A., Regev, I., Mahadevan, L., and Pourquié, O. (2017). Excitable Dynamics and Yap-Dependent Mechanical Cues Drive the Segmentation Clock. Cell 171, 668–682.e11. 10.1016/j.cell.2017.08.043.

19. Cooke, J., and Zeeman, E.C. (1976). A clock and wavefront model for control of the number of repeated structures during animal morphogenesis. J. Theor. Biol. 58, 455–476. 10.1016/s0022-5193(76)80131-2.

20. Dubrulle, J., McGrew, M.J., and Pourquié, O. (2001). FGF Signaling Controls Somite Boundary Position and Regulates Segmentation Clock Control of Spatiotemporal Hox Gene Activation. Cell 106, 219–232. 10.1016/s0092-8674(01)00437-8.

21. Durbin, L., Brennan, C., Shiomi, K., Cooke, J., Barrios, A., Shanmugalingam, S., Guthrie, B., Lindberg, R., and Holder, N. (1998). Eph signaling is required for segmentation and differentiation of the somites. Genes Dev. 12, 3096–3109. 10.1101/gad.12.19.3096.

22. Nakaya, Y., Kuroda, S., Katagiri, Y.T., Kaibuchi, K., and Takahashi, Y. (2004). Mesenchymal-Epithelial Transition during Somitic Segmentation Is Regulated by Differential Roles of Cdc42 and Rac1. Dev. Cell 7, 425–438. 10.1016/j.devcel.2004.08.003.

23. Watanabe, T., Sato, Y., Saito, D., Tadokoro, R., and Takahashi, Y. (2009). EphrinB2 coordinates the formation of a morphological boundary and cell epithelialization during somite segmentation. Proc. Natl. Acad. Sci. 106, 7467–7472. 10.1073/pnas.0902859106.

24. Dahmann, C., Oates, A.C., and Brand, M. (2011). Boundary formation and maintenance in tissue development. Nat Rev Genet 12, 43–55. 10.1038/nrg2902.

25. Julich, D., Mould, A.P., Koper, E., and Holley, S.A. (2009). Control of extracellular matrix assembly along tissue boundaries via Integrin and Eph/Ephrin signaling. Development 136, 2913–2921. 10.1242/dev.038935.

26. Lackner, S., Schwendinger-Schreck, J., Jülich, D., and Holley, S.A. (2013). Segmental Assembly of Fibronectin Matrix Requires rap1b and integrin α5. Dev. Dyn. 242, 122–131. 10.1002/dvdy.23909.

27. McMillen, P., Chatti, V., Jülich, D., and Holley, S.A. (2016). A sawtooth pattern of cadherin 2 stability mechanically regulates somite morphogenesis. Curr Biol 26, 542–549. 10.1016/j.cub.2015.12.055.

28. Naganathan, S.R., and Oates, A.C. (2020). Patterning and mechanics of somite boundaries in zebrafish embryos. Semin. Cell Dev. Biol. 107, 170–178. 10.1016/j.semcdb.2020.04.014.

29. Tlili, S., Yin, J., Rupprecht, J.-F., Mendieta-Serrano, M.A., Weissbart, G., Verma, N., Teng, X., Toyama, Y., Prost, J., and Saunders, T.E. (2019). Shaping the zebrafish myotome by intertissue friction and active stress. Proc. Natl. Acad. Sci. 116, 25430–25439. 10.1073/pnas.1900819116.

30. Bard, J.B.L. (1988). A traction-based mechanism for somitogenesis in the chick. Roux’s Arch. Dev. Biol. 197, 513–517. 10.1007/bf00385686.

31. Packard, D.S., and Jacobson, A.G. (1979). Analysis of the physical forces that influence the shape of chick somites. J. Exp. Zoöl. 207, 81–92. 10.1002/jez.1402070109.

32. Dias, A.S., Almeida, I. de, Belmonte, J.M., Glazier, J.A., and Stern, C.D. (2014). Somites Without a Clock. Science 343, 791–795. 10.1126/science.1247575.

33. Nelemans, B.K.A., Schmitz, M., Tahir, H., Merks, R.M.H., and Smit, T.H. (2020). Somite Division and New Boundary Formation by Mechanical Strain. iScience 23, 100976. 10.1016/j.isci.2020.100976.

34. Mongera, A., Rowghanian, P., Gustafson, H.J., Shelton, E., Kealhofer, D.A., Carn, E.K., Serwane, F., Lucio, A.A., Giammona, J., and Campàs, O. (2018). A fluid-to-solid jamming transition underlies vertebrate body axis elongation. Nature 561, 401–405. 10.1038/s41586-018-0479-2.

35. Serwane, F., Mongera, A., Rowghanian, P., Kealhofer, D.A., Lucio, A.A., Hockenbery, Z.M., and Campàs, O. (2017). In vivo quantification of spatially varying mechanical properties in developing tissues. Nat Methods 14, 181–186. 10.1038/nmeth.4101.

36. Shelton, E., Serwane, F., and Campas, O. (2017). Geometrical characterization of fluorescently labelled surfaces from noisy 3D microscopy data. Journal of microscopy 269, 259–268. 10.1111/jmi.12624.

37. Gross, B.J., Soltwedel, J.R., Shelton, E., Gomez, C., and Campàs, O. (2025). STRESS, an automated geometrical characterization of deformable particles for in vivo measurements of cell and tissue mechanical stresses. Sci. Rep. 15, 28599. 10.1038/s41598-025-13419-z.

38. Mongera, A., Pochitaloff, M., Gustafson, H.J., Stooke-Vaughan, G.A., Rowghanian, P., Kim, S., and Campàs, O. (2023). Mechanics of the cellular microenvironment as probed by cells in vivo during zebrafish presomitic mesoderm differentiation. Nat Mater 22, 135–143. 10.1038/s41563-022-01433-9.

39. Kim, S., Pochitaloff, M., Stooke-Vaughan, G.A., and Campàs, O. (2021). Embryonic tissues as active foams. Nat Phys 17, 859–866. 10.1038/s41567-021-01215-1.

40. Umetsu, D., Aigouy, B., Aliee, M., Sui, L., Eaton, S., Jülicher, F., and Dahmann, C. (2014). Local Increases in Mechanical Tension Shape Compartment Boundaries by Biasing Cell Intercalations. Curr. Biol. 24, 1798–1805. 10.1016/j.cub.2014.06.052.

41. Landsberg, K.P., Farhadifar, R., Ranft, J., Umetsu, D., Widmann, T.J., Bittig, T., Said, A., Jülicher, F., and Dahmann, C. (2009). Increased Cell Bond Tension Governs Cell Sorting at the Drosophila Anteroposterior Compartment Boundary. Curr Biol 19, 1950–1955. 10.1016/j.cub.2009.10.021.

42. Monier, B., Pélissier-Monier, A., Brand, A.H., and Sanson, B. (2010). An actomyosin-based barrier inhibits cell mixing at compartmental boundaries in Drosophila embryos. Nat. Cell Biol. 12, 60–65. 10.1038/ncb2005.

43. Major, R.J., and Irvine, K.D. (2006). Localization and requirement for Myosin II at the dorsal-ventral compartment boundary of the Drosophila wing. Dev. Dyn. 235, 3051–3058. 10.1002/dvdy.20966.

44. Koshida, S., Kishimoto, Y., Ustumi, H., Shimizu, T., Furutani-Seiki, M., Kondoh, H., and Takada, S. (2005). Integrinα5-Dependent Fibronectin Accumulation for Maintenance of Somite Boundaries in Zebrafish Embryos. Developmental Cell 8, 587–598. 10.1016/j.devcel.2005.03.006.

45. Barrios, A., Poole, R.J., Durbin, L., Brennan, C., Holder, N., and Wilson, S.W. (2003). Eph/Ephrin Signaling Regulates the Mesenchymal-to-Epithelial Transition of the Paraxial Mesoderm during Somite Morphogenesis. Curr. Biol. 13, 1571–1582. 10.1016/j.cub.2003.08.030.

46. Fiore, A., Yu, G., Northey, J.J., Patel, R., Ravenscroft, T.A., Ikegami, R., Kolkman, W., Kumar, P., Dilan, T.L., Ruetten, V.M.S., et al. (2025). Live imaging of the extracellular matrix with a glycan-binding fluorophore. Nat. Methods 22, 1070–1080. 10.1038/s41592-024-02590-2.

47. Lele, Z., Folchert, A., Concha, M., Rauch, G.-J., Geisler, R., Rosa, F., Wilson, S.W., Hammerschmidt, M., and Bally-Cuif, L. (2002). parachute/n-cadherin is required for morphogenesis and maintained integrity of the zebrafish neural tube. Development 129, 3281– 3294.

48. Naganathan, S.R., Popović, M., and Oates, A.C. (2022). Left–right symmetry of zebrafish embryos requires somite surface tension. Nature, 1–6. 10.1038/s41586-022-04646-9.

49. Yanagida, A., Corujo-Simon, E., Revell, C.K., Sahu, P., Stirparo, G.G., Aspalter, I.M., Winkel, A.K., Peters, R., Belly, H.D., Cassani, D.A.D., et al. (2022). Cell surface fluctuations regulate early embryonic lineage sorting. Cell 185, 777–793.e20. 10.1016/j.cell.2022.01.022.

50. Cayuso, J., Xu, Q., Addison, M., and Wilkinson, D.G. (2019). Actomyosin regulation by Eph receptor signaling couples boundary cell formation to border sharpness. eLife 8, e49696. 10.7554/elife.49696.

51. Cayuso, J., Xu, Q., and Wilkinson, D.G. (2015). Mechanisms of boundary formation by Eph receptor and ephrin signaling. Dev. Biol. 401, 122–131. 10.1016/j.ydbio.2014.11.013.

52. Kindberg, A.A., Srivastava, V., Muncie, J.M., Weaver, V.M., Gartner, Z.J., and Bush, J.O. (2021). EPH/EPHRIN regulates cellular organization by actomyosin contractility effects on cell contacts. J. Cell Biol. 220, e202005216. 10.1083/jcb.202005216.

53. Jülich, D., Cobb, G., Melo, A.M., McMillen, P., Lawton, A.K., Mochrie, S.G.J., Rhoades, E., and Holley, S.A. (2015). Cross-Scale Integrin Regulation Organizes ECM and Tissue Topology. Dev Cell 34, 33–44. 10.1016/j.devcel.2015.05.005.

54. Jülich, D., and Holley, S.A. (2024). Live imaging of Fibronectin 1a-mNeonGreen and Fibronectin 1b-mCherry knock-in alleles during early zebrafish development. Cells Dev. 177, 203900. 10.1016/j.cdev.2024.203900.

55. Veenvliet, J.V., Bolondi, A., Kretzmer, H., Haut, L., Scholze-Wittler, M., Schifferl, D., Koch, F., Guignard, L., Kumar, A.S., Pustet, M., et al. (2020). Mouse embryonic stem cells self-organize into trunk-like structures with neural tube and somites. Science 370. 10.1126/science.aba4937.

56. Kulesa, P.M., and Fraser, S.E. (2002). Cell Dynamics During Somite Boundary Formation Revealed by Time-Lapse Analysis. Science 298, 991–995. 10.1126/science.1075544.

57. Nüsslein-Volhard, C., and Dahm, R. (2002). Zebrafish (Oxford University Press).

58. Cooper, M.S., Szeto, D.P., Sommers-Herivel, G., Topczewski, J., Solnica-Krezel, L., Kang, H., Johnson, I., and Kimelman, D. (2005). Visualizing morphogenesis in transgenic zebrafish embryos using BODIPY TR methyl ester dye as a vital counterstain for GFP. Dev. Dyn. 232, 359–368. 10.1002/dvdy.20252.

59. Swinburne, I.A., Mosaliganti, K.R., Upadhyayula, S., Liu, T.-L., Hildebrand, D.G.C., Tsai, T.Y.-C., Chen, A., Al-Obeidi, E., Fass, A.K., Malhotra, S., et al. (2018). Lamellar projections in the endolymphatic sac act as a relief valve to regulate inner ear pressure. eLife 7, e37131. 10.7554/elife.37131.

60. Compagnon, J., Barone, V., Rajshekar, S., Kottmeier, R., Pranjic-Ferscha, K., Behrndt, M., and Heisenberg, C.-P. (2014). The Notochord Breaks Bilateral Symmetry by Controlling Cell Shapes in the Zebrafish Laterality Organ. Dev. Cell 31, 774–783. 10.1016/j.devcel.2014.11.003.

61. Maitre, J.L., Berthoumieux, H., Krens, S.F.G., Salbreux, G., Julicher, F., Paluch, E., and Heisenberg, C.P. (2012). Adhesion Functions in Cell Sorting by Mechanically Coupling the Cortices of Adhering Cells. Science 338, 253–256. 10.1126/science.1225399.

62. Revenu, C., Streichan, S., Dona, E., Lecaudey, V., Hufnagel, L., and Gilmour, D. (2014). Quantitative cell polarity imaging defines leader-to-follower transitions during collective migration and the key role of microtubule-dependent adherens junction formation. Development 141, 1282– 1291. 10.1242/dev.101675.

63. Goody, M.F., Kelly, M.W., Lessard, K.N., Khalil, A., and Henry, C.A. (2010). Nrk2b-mediated NAD+ production regulates cell adhesion and is required for muscle morphogenesis in vivo Nrk2b and NAD+ in muscle morphogenesis. Dev. Biol. 344, 809–826. 10.1016/j.ydbio.2010.05.513.

64. Soans, K.G., Ramos, A.P., Sidhaye, J., Krishna, A., Solomatina, A., Hoffmann, K.B., Schlüßler, R., Guck, J., Sbalzarini, I.F., Modes, C.D., et al. (2022). Collective cell migration during optic cup formation features changing cell-matrix interactions linked to matrix topology. Curr. Biol. 32, 4817–4831.e9. 10.1016/j.cub.2022.09.034.

65. Lele, Z., Folchert, A., Concha, M., Rauch, G.-J., Geisler, R., Rosa, F., Wilson, S.W., Hammerschmidt, M., and Bally-Cuif, L. (2002). parachute / n-cadherin is required for morphogenesis and maintained integrity of the zebrafish neural tube. Development 129, 3281– 3294. 10.1242/dev.129.14.3281.

66. LaMora, A., and Voigt, M.M. (2009). Cranial sensory ganglia neurons require intrinsic N-cadherin function for guidance of afferent fibers to their final targets. Neuroscience 159, 1175– 1184. 10.1016/j.neuroscience.2009.01.049.

67. Jülich, D., Geisler, R., Consortium, T. 2000 S., and Holley, S.A. (2005). Integrinα5 and Delta/Notch Signaling Have Complementary Spatiotemporal Requirements during Zebrafish Somitogenesis. Dev. Cell 8, 575–586. 10.1016/j.devcel.2005.01.016.

68. Schindelin, J., Arganda-Carreras, I., Frise, E., Kaynig, V., Longair, M., Pietzsch, T., Preibisch, S., Rueden, C., Saalfeld, S., Schmid, B., et al. (2012). Fiji: an open-source platform for biological-image analysis. Nat Methods 9, 676–682. 10.1038/nmeth.2019.

69. Pourquié, O., and Tam, P.P.L. (2001). A Nomenclature for Prospective Somites and Phases of Cyclic Gene Expression in the Presomitic Mesoderm. Dev. Cell 1, 619–620. 10.1016/s1534-5807(01)00082-x.

70. Schröter, C., Herrgen, L., Cardona, A., Brouhard, G.J., Feldman, B., and Oates, A.C. (2008). Dynamics of zebrafish somitogenesis. Dev. Dyn. 237, 545–553. 10.1002/dvdy.21458.

71. Holtze, C., Rowat, A.C., Agresti, J.J., Hutchison, J.B., Angile, F.E., Schmitz, C.H.J., Koster, S., Duan, H., Humphry, K.J., Scanga, R.A., et al. (2008). Biocompatible surfactants for water-in-fluorocarbon emulsions. Lab Chip 8, 1632–1639. 10.1039/b806706f.

72. Lim, I., Vian, A., Wouw, H.L. van de, Day, R.A., Gomez, C., Liu, Y., Rheingold, A.L., Campàs, O., and Sletten, E.M. (2020). Fluorous Soluble Cyanine Dyes for Visualizing Perfluorocarbons in Living Systems. J Am Chem Soc 142, 16072–16081. 10.1021/jacs.0c07761.

73. Campas, O., Mammoto, T., Hasso, S., Sperling, R.A., O’Connell, D., Bischof, A.G., Maas, R., Weitz, D.A., Mahadevan, L., and Ingber, D.E. (2014). Quantifying cell-generated mechanical forces within living embryonic tissues. Nat Methods 11, 183–189. 10.1038/nmeth.2761.

74. Krull, A., Buchholz, T.-O., and Jug, F. (2019). Noise2Void - Learning Denoising from Single Noisy Images. 2019 IEEECVF Conf. Comput. Vis. Pattern Recognit. (CVPR) 00, 2124–2132. 10.1109/cvpr.2019.00223.

75. Boutillon, A., Escot, S., Elouin, A., Jahn, D., González-Tirado, S., Starruß, J., Brusch, L., and David, N.B. (2022). Guidance by followers ensures long-range coordination of cell migration through α-catenin mechanoperception. Dev. Cell 57, 1529–1544.e5. 10.1016/j.devcel.2022.05.001.

76. Boutillon, A., Escot, S., and David, N.B. (2021). Deep and Spatially Controlled Volume Ablations using a Two-Photon Microscope in the Zebrafish Gastrula. J. Vis. Exp. 10.3791/62815.

77. Aigouy, B., Umetsu, D., and Eaton, S. (2016). Drosophila, Methods and Protocols. Methods Mol. Biol. 1478, 227–239. 10.1007/978-1-4939-6371-3_13.

78. Guntas, G., Hallett, R.A., Zimmerman, S.P., Williams, T., Yumerefendi, H., Bear, J.E., and Kuhlman, B. (2015). Engineering an improved light-induced dimer (iLID) for controlling the localization and activity of signaling proteins. Proc. Natl. Acad. Sci. 112, 112–117. 10.1073/pnas.1417910112.

79. Guo, H., Swan, M., and He, B. (2022). Optogenetic inhibition of actomyosin reveals mechanical bistability of the mesoderm epithelium during Drosophila mesoderm invagination. eLife 11, e69082. 10.7554/elife.69082.

80. Yamamoto, K., Miura, H., Ishida, M., Mii, Y., Kinoshita, N., Takada, S., Ueno, N., Sawai, S., Kondo, Y., and Aoki, K. (2021). Optogenetic relaxation of actomyosin contractility uncovers mechanistic roles of cortical tension during cytokinesis. Nat. Commun. 12, 7145. 10.1038/s41467-021-27458-3.

